# Kir6.1, a component of an ATP-sensitive potassium channel, regulates natural killer cell development

**DOI:** 10.1101/2024.08.14.608003

**Authors:** Natalie Samper, Lilja Harðardóttir, Delphine M Depierreux, Soomin C. Song, Ayano Nakazawa, Ivan Gando, Tomoe Y. Nakamura, Andrew M Sharkey, Carla R. Nowosad, Stefan Feske, Francesco Colucci, William A. Coetzee

**Author notes:** Corresponding author: Dr. William A. Coetzee, NYU School of Medicine, 550 1st Avenue, Smilow 604, New York, NY 10016, USA, Corresponding author: Dr. Francesco Colucci, Professor of Immunology, Clinical School, University of Cambridge, King’s College, Cambridge, UK. These authors have contributed equally (Natalie Samper, Lilja Harðardóttir, Delphine M Depierreux).

## Abstract

Involved in immunity and reproduction, natural killer (NK) cells offer opportunities to develop new immunotherapies to treat infections and cancer or to alleviate pregnancy complications. Most current strategies use cytokines or antibodies to enhance NK-cell function, but none use ion channel modulators, which are widely used in clinical practice to treat hypertension, diabetes, epilepsy, and other conditions. Little is known about ion channels in NK cells. We show that *Kcnj8,* which codes for the Kir6.1 subunit of a certain type of ATP-sensitive potassium (K_ATP_) channel, is highly expressed in murine splenic and uterine NK cells compared to other K^+^ channels previously identified in NK cells. *Kcnj8* expression is highest in the most mature subset of splenic NK cells (CD27^-^CD11b^+^) and in NKG2A^+^ or Ly49C/I^+^ educated uterine NK cells. Using patch clamping, we show that a subset of NK cells expresses a current sensitive to the Kir6.1 blocker PNU-37883A. *Kcnj8* does not participate in NK cell degranulation in response to tumor cells *in vitro* or rejection of tumor cells *in vivo*. Transcriptomics show that genes previously implicated in NK cell development are amongst those differentially expressed in CD27^-^CD11b^+^ NK cells deficient of *Kcnj8*. Indeed, we found that mice with NK-cell specific *Kcnj8* gene ablation have fewer CD11b^+^CD27^-^ and KLRG-1^+^ NK cells in the bone barrow and spleen. These results show that the K_ATP_ subunit Kir6.1 has a key role in NK-cell development.

## Introduction

Natural Killer (NK) cells have a key role in immune surveillance, anti-tumor immunity and anti-viral immunity by identifying and eliminating transformed or infected cells. NK cells lyse these target cells through a process of regulated exocytosis of specialized secretory lysosomes, termed cytotoxic granules, which contain proteins such as perforin, granzymes, and the Fas ligand. A second major effector function of NK cells is to release chemokines and cytokines, which occurs after binding to a target cell, or when NK cells are activated by cytokines and chemokines secreted from other immune cells. The cytokines released from NK cells, including interferon-γ (IFN-γ), provide feedback to other immune cell types and modulate the innate and adaptive immune function. IFN-γ is a pleiotropic cytokine that is critical for cancer immune surveillance (1) by directly interacting with cancer cells to reduce cell proliferation and induce apoptosis. Thus, cytokine and chemokine secretion and perforin-mediated cytotoxicity are two independent effector functions that largely account for the antiviral and antitumor activities of NK cells. A special type of NK cell in the uterus (uNK) mediates physiological vascular changes that enable the placenta to sustain healthy fetal growth (2). Understanding the nuances of NK cell development, education and activation is essential for elucidating the mechanisms underlying NK cell function and for developing therapeutic strategies targeting NK cells in various disease settings.

NK cells develop in the bone marrow where integrins and other surface markers are acquired. In C57BL/6 mice, the maturation program is marked by sequential surface expression of integrin subunit integrin alpha M (*Itgam*), also known as CD11b, MAC-1 or CR3, and the TNF-superfamily CD27 receptor (3). CD27^−^CD11b^−^ double negative (DN) cells develop in the bone marrow and are the most immature. After the acquisition of CD27 and CD11b, expression of S1P5R enables the CD27^+^CD11b^+^ double-positive (DP) NK cells to migrate from the bone marrow to the periphery, including the blood, spleen, and other organs. With further maturation, CD27 expression is lost and CD27^−^CD11b^+^ remains as the most mature population with the highest cytotoxic potential.

In both mice and humans, NK cells possess inhibitory receptors that recognize self-MHC class I molecules. These receptors include killer cell immunoglobulin-like receptors (KIRs) in humans, Ly49 receptors in mice, and CD94/NKG2A in both species (4). Although some NK-cell receptors differ between humans and different mouse strains, the intracellular signals that govern NK cell activation are remarkably conserved (5). Biochemical signals downstream of inhibitory and activating receptors are transduced inside NK cells by receptor-associated tyrosine phosphatases and kinases, e.g. SHP-1, which regulate Vav-1, PLC-γ2, and other key intracellular signaling molecules that mediate degranulation (6). NK-cell education (also referred to as licensing) occurs through homeostatic interactions between self MHC class I molecules and NK cell inhibitory receptors (*7*), ensuring that NK cells effectively distinguish between healthy cells and those with inadequate MHC class I expression due to infection or transformation. When NK cells interact with cells expressing self-MHC class I molecules, inhibitory signals are transmitted through these receptors, preventing NK cell activation and cytotoxicity against healthy cells. NK cells can directly recognize cells with aberrant expression of MHC class I molecules (“missing self”) or cells expressing stress-induced ligands or viral proteins (“induced self”). Human MIC-A/B or mouse Rae1, for example, engage the activating receptor NKG2D in both species. Engagement of activating receptors in the absence of ligands for inhibitory receptors initiates signaling pathways leading to NK cell activation. NK cell activation can also be triggered by cytokines such as interleukin-2 (IL-2), IL-12, IL-15, IL-18, and interferon-alpha (IFN-α), the latter induced experimentally in mice by administration of the viral RNA mimic polyinosinic:polycytidylic acid (poly I:C). Once activated, NK cells release cytotoxic perforin and granzyme molecules and produce inflammatory cytokines, such as interferon-γ (IFN-γ).

Ion translocating proteins regulate immune cell function. There are hundreds such proteins that control the movement of ions such as Na^+^, K^+^, Cl^-^, Ca^2+^, Mg^2+^, Zn^2+^ across the plasma membrane. Although extensively studied in the nervous, endocrine, and cardiovascular systems, the roles of this important class of proteins are scantly studied in immune cells (8). In T cells, B cells, monocytes, macrophages, dendritic cells and neutrophils, roles have been described for certain K^+^ channels (KCa3.1 and Kv1.3), Ca^2+^ channels (mainly ORAI1/STIM1), P2X receptors, and some members of the transient receptor potential (TRP) family (9). The involvement of some of these channels in immunity is highlighted by inherited gene variants (channelopathies) in immunodeficiency disorders. By contrast, ion channels, exchangers and pumps in NK cells are poorly characterized. There are only a few studies describing roles of ion channels in human blood peripheral NK cells. Studies in the 1980’s have identified that the K^+^ concentration and some ion channel blockers can regulate NK cell-mediated killing (10–12). A sprinkling of studies described the existence of subtypes of ion channels in NK cells, including voltage-gated K^+^ channels (13, 14), Ca^2+^-activated K^+^ channels (15, 16), two-pore domain K^+^ channels (17), Ca^2+^-permeable cation channels (18), intracellular Ca^2+^-permeable channels (19, 20), and store-operated Ca^2+^ entry (SOCE) channels (21). The lack of a complete understanding of the role of ion translocating mechanisms in NK cells is a major gap in our knowledge of immunity to infection, tumors, autoimmunity, and inflammation. This is also a possible missed opportunity for the development of novel therapeutic approaches in inflammatory and anti-cancer therapy.

ATP-sensitive K^+^ (K_ATP_) channels are regulated by intracellular nucleotides (ATP and ADP). As such, they couple the intracellular energy metabolic state to membrane excitability and secretory events (22, 23). In the pancreatic β-cell, for example, K_ATP_ channel activity initiates the release of insulin from intracellular granules after a meal (24), whereas in the heart they participate in stress responses (22). The K_ATP_ channel is a tetrameric structure composed of four pore-forming subunits (Kir6.1 or Kir6.2) in association with four sulphonylurea receptors (SUR1 or SUR2) (25, 26) (**Figure S1**). *KCNJ8* and *ABCC9* are genes adjacent to each other on human chromosome 12 and respectively code for Kir6.1 and SUR2 subunits, whereas the adjacent *KCNJ11* and *ABCC8* on chromosome 11 respectively code for Kir6.2 and SUR1 subunits. The aim of this study was to characterize the expression of *Kcnj8* in mouse NK cells and to determine whether *Kcnj8* has a role in NK cell development and function. We found *Kcnj8* to be expressed at high levels in a mature subset of mouse NK cells. NK-cell specific *Kcnj8* gene ablation did not interfere with the basic NK-cell functions that we studied, but impeded full NK cell maturation.

## Methods

### Generation of NK cell-specific Kir6.1 knockout mice

The mouse model was generated by inGenious Targeting Labs Inc. (iTL; Stony Brook, New York). In brief, a 10.4 kb region used to construct the targeting vector was first subcloned from a positively identified C57BL/B6 BAC clone (RPCI-23: 108N15) BAC clone. The region was designed such that the short homology arm (SA) extended ∼1.76 kb 3’ to exon 2. The ∼7.2 kb long homology arm (LA) ends 5’ to exon 2. The loxP/FRT flanked Neo cassette was inserted on the 3’ side of exon 2 and the single loxP site is inserted on the 5’ of exon 2. The target region was 1,460 bps and includes exon 2. The targeting vector was linearized with NotI and transfected by electroporation of BA1 (C57BL/6 x 129/SvEv) hybrid embryonic stem cells. After selection with G418 antibiotic, surviving clones were expanded for PCR analysis to identify recombinant ES clones. Hybrid positive ES cells were injected into C57BL/6 blastocysts. The Neo cassette was removed by breeding to FLP deleter mice (Jackson stock #005703), followed by backcrossing to a C57BL/6 background for >10 generations to achieve congenicity. To target NK cells, we used inducible NKp46-CreERT2 mice from Dr. Lewis Lanier (UC San Francisco, USA) (27) that were intercrossed with Rosa26-tdTomato C57BL/6 mice. The Kir6.1^flx/flx^ mice were crossed with NKp46-CreERT2 and bred to homozygosity. Kir6.1 was deleted in NK cells by treating mice with tamoxifen (IP injection of 2 mg tamoxifen in corn oil, daily for 4 days). In some experiments we used a similar mouse model in which exon 2 of *Kcnj8* is flanked by loxP sites (provided by Dr. Andrew Tinker) that was crossed with constitutively expressed NKp46-Cre mice provided by Dr. Eric Vivier (Aix-Marseille University, France) (28).

### Isolation of mouse splenic NK cells

All animal handling procedures were in accordance with National Institutes of Health guidelines and were approved by the Institutional Animal Care and Use Committees of New York University School of Medicine and the University of Cambridge. Male and female C57BL/6 mice aged 8-12 weeks were housed in pathogen-free conditions, before being euthanized and collection of organs such as the spleen, brain or bone marrow. For bone marrow cells, the femur was collected, cells were flushed with PBS in a syringe and passed through a 70 µm cell strainer followed by washing with PBS. Cells from peritoneal cavity were aspirated with a syringe and washed with PBS. In some experiments, spleens were harvested in media (RPMI + 10% FBS) on ice and passed through 70 µm cell strainer along with media and centrifuged. Erythrocytes were lysed with 1xRBC lysis buffer (Ebioscience) for 3 min at RT. Splenocytes were washed 2x with media and passed again through 70 µm cell strainer. In other experiments, spleens were dissociated with the gentleMACS spleen disassociation kit (Miltenyi Biotec). Cells were then filtered, erythrocytes were depleted via isotonic hemolysis, and NK cells were isolated using a negative selection method (NK Cell Isolation Kit, Miltenyi Biotec) following the manufacture’s guidelines. In this negative selection method, non-NK cells, i.e. T cells, dendritic cells, B cells, granulocytes, macrophages, and erythroid cells are magnetically labelled with a cocktail of biotin-conjugated antibodies and anti-biotin MicroBeads, and then removed from isolated splenic cells using magnetic columns.

### Single cell RNA seq of splenic NK cells

Splenic NK cells from 3 wildtype mice were isolated via negative selection as previously described. Cells were lysed and RNA was obtained via PicoPure™ RNA Isolation Kit (Applied Biosystems). Lipid-based cell multiplexing was used in order to combine NK cell populations isolated from 3 mice. Single cell RNA sequencing (RNA seq) was coordinated by NYU Langone’s Genome Technology Center (RRID: SCR_017929). We used the Chromium Next GEM Single Cell 3ʹ Reagent Kit v3.1 (10x Genomics) to perform gel beads-in-emulsion (GEM) generation and barcoding, Post GEM-reverse transcriptase (RT) cleanup and cDNA amplification, and 3’ gene expression library construction. Sequencing was performed by NYU Langone’s Genome Technology Center using an Illumina NovaSeq 6000 instrument. Demultiplexing raw base call (BCL) files into FASTQ format was performed using Cell Ranger software (10X Genomics). The latter software was also used for alignment, filtering, barcode counting, and UMI counting to generate feature-barcode matrices. The estimated number of cells was 16,384, with 72,726 mean reads per cell (2,047 median reads per cell). Of a total of 1,191,540,284 reads, 96.6% was mapped to the mm10 mouse genome.

Further analysis including quality filtering, the identification of highly variable genes, dimensionality reduction, standard unsupervised clustering algorithms, and the discovery of differentially expressed genes was performed using the Seurat R package (29). We filtered out cells with unique feature counts (genes) over 5,000 or less than 200. We also filtered out cells with more than 15% mitochondrial gene counts. Global-scaling normalization (“LogNormalize”) was performed that normalizes the feature expression measurements for each cell by the total expression, multiplies this by a scale factor (10,000) and log-transforms the result. Linear transformation (scaling) was performed, followed by dimensional reduction using a principal component analysis (PCA). Nearest-neighbor analysis, constrained to a maximum of 20 dimensions, was performed, followed by a graph-based approach to cluster cells at a resolution value of 0.5. Clusters were displayed after non-linear dimensional reduction with t-distributed stochastic neighbor embedding (tSNE) and uniform manifold approximation and projection (UMAP) methods. Differentially overexpressed features (cluster biomarkers) were determined using the ‘FindAllMarkers’ function at a minimum fold difference (log-scale) of 0.25.

### Bulk RNA seq of mouse splenic NK cells

Splenocytes were isolated from wild-type mice, and Kir6.1^flx/flx^ x NKp46-CreERT2 mice immediately following 4 days of tamoxifen induction (n=2 each). Cells were first gated for NK cell markers NK1.1 and NKp46, then sorted into four pools (CD27^-^/CD27b^-^, CD27^+^/CD27b^-^, CD27^+^/CD27b^+^, and CD27^-^/CD27b^+^). RNA was isolated with the RNeasy Mini Kit (Qiagen). RNA extractions were quantified using RNA Pico Chips (Agilent, Cat. 5067-1513) on an Agilent 2100 BioAnalyzer. cDNA was synthesized using the Takara SMART-Seq HT kit (Cat. 634438) with 0.5 ng of RNA input and 14 amplification cycles. The Nextera XT DNA library Kit (Illumina, Cat. FC-131-1096) was used to construct sequencing libraries with 0.25 ng cDNA input and 11 cycles amplification. Final libraries were visualized using High Sensitivity DNA ScreenTape (Agilent, Cat. #5067-5584) on the Agilent Tapestation 2200 instrument. Quant-It (Invitrogen, Cat. P11495) was used for final concentration determination and libraries were pooled equimolar. Paired-end 50 cycle RNA sequencing was performed by NYU Langone’s Genome Technology Center (RRID: SCR_017929) using a single lane of a 10B 100 Cycle Flowcell using an Illumina NovaSeq X-Plus instrument. Data analysis was performed on BigPurple, a high-performance computing cluster available through NYU Langone’s High Performance Computing (HPC) Core. Quality control of the paired RNA fastq files was performed with fastqc. No end-trimming was needed. Reads were aligned to the mm10 mouse genome using STAR and index files were produced with Samtools. The GeneCounts option of STAR was used to count number reads per gene while mapping. Off-line analysis was performed with R. Individual “ReadsPerGene.out.tab” files were combined. Gene expression analysis was performed with the DESeq2 package (version 1.44.0), which included filtering low expressing genes, calculating normalized gene counts, calculating the between-sample distance matrix, principal component analysis, and performing a differential expression analysis. Gene ontology analysis was performed with Enrichr (30). The scripts used for analysis are available upon request.

### Bulk RNA seq of uNK cells

At gestation day 9.5-10.5, C57BL6 mice were culled by cervical dislocation to collect the pregnant uterus which was enzymatically and mechanically processed into a single suspension as previously described (31). Single cell suspensions were labelled with viability dye Zombie Aqua Fixable Viability Kit according to the manufacturer’s protocol. Fc receptors were blocked with anti-CD16/32 mAb (Trustain Fcx) for 10 min at 4 °C. Cell surface antigens were labelled for 30 min at 4 °C in the dark, with the indicated cocktail of antibodies (**Table S1**). Cells were subsequently washed twice in PBS, and collected by centrifugation at 400 *g* for 5 min at 4 °C. Live single cells, negative for CD3, CD4, CD8, CD19, CD11b high, and positive for CD45 and Nkp46 were defined as NK cells. NK cells were further gated according to their relative expression of LY49C/I and NKG2A. Three cell populations were collected (Ly49C/I neg, NKG2A neg ; LY49C/I neg, NKG2A pos ; LY49C/I pos NKG2A neg). Double positive cells were not collected due to limited sample size. Samples were stored at −80 °C until RNA extraction. RNA procedures and library construction was performed as previously described (32). Reads were trimmed using TrimGalore v0.5.0. and were mapped using STAR v2.7.1. to the Ensembl Mus_musculus GRCm38 (release 103) reference genome. Differential gene expression analysis was done using the counted reads and the R package edgeR version 3.26.5 (R version 3.6.1). The threshold was set to include genes with at least 5 counts per million reads mapped and detected in at least half the samples to allow for on/off expression pattern detection. Correction for GC content and gene length bias was applied using CQN bioconductor package (version 1.30.0). The resulting p-value was corrected for multiple hypothesis testing using the Benjamini-Hochberg method.

### qRT-PCR experiments

Splenic NK cells isolated with a negative selection method were used. First strand cDNA synthesis was performed with SuperScript™ IV (ThermoFisher Scientific) using random primers. Primers used for PCR spanned the boundary of exons 1 and 2 of *Kcnj8* (1585F: 5’-CACAAGAACATCCGAGAGC-3’ and 1585R: 5’-GGGCATTCCTCAGTCATCAT-3’), which produced a 320bp amplicon. The cycling conditions consisted of heating to 95 °C for 3 min, 28 cycles consisting of 94 °C for 30s, 55 °C for 45s, 72 °C for 1 min; followed by 1 min extension at 72 °C. Amplicons were resolved by 1% agarose gel electrophoresis and visualized with ethidium bromide.

### Flow cytometry

Single cell suspensions were obtained from organs. Briefly, cells were blocked with TruStain FcX™ (anti-mouse CD16/32) antibody (Biolegend) according to the manufacturer’s protocol. Cells were stained with relevant antibodies for 30 min on ice in the dark followed by two washes with PBS. When detecting intracellular antigens, the cells were fixed and permeabilized with eBioscience Fixation/Permeabilization kit, according to manufacturer’s protocol prior to staining with intracellular antibodies. Data was acquired on a Cytek Aurora instrument.

Antibodies used for flow cytometry: Fixable Viability Dye eFluor™ 780 (eBioscience), CD3 BV785 (Biolegend), NK1.1 PE (Biolegend), CD11b BV421 (BD Biosciences), CD27 BV605 (BD Biosciences), CD49b Pe-Cy7 (Biolegend), DNAM1 AF647 (Biolegend), KLRG1 FITC (eBioscience), h2-kb BV421 (Biolegend), CD107a FITC (BD Biosciences).

In some experiments, either whole spleen or isolated NK cells were incubated in PEB buffer (PBS+0.5% BSA+2mM EDTA) with TruStain FcX™ (anti-mouse CD16/32) antibody (Biolegend) on ice for 5 minutes. Cells were then stained with the appropriate conjugated surface markers in PEB buffer protected from light at 4 °C for 20 min. In the case of intracellular staining, cells were incubated with IC Fixation Buffer (eBioscience) for 30 min and then Permeabilization Buffer (eBioscience) with addition of conjugated antibody overnight. Anti-mouse antibodies used cytometry: NKp46 (29A1.4) BV421 (Biolegend), NK1.1 (PK136) APC-Cy7 (Biolegend), NK1.1 (PK136) PE (Biolegend), CD11b (M1/70) APC (Biolegend), CD11b (M1/70) AF488 (Biolegend), CD27 (LG.3A10) APC-Cy7 (Biolegend), CD27 (LG.3A10) AF647, and CD107a (1D4B) APC/Fire750 (Biolegend).

### Degranulation assay with target cells

In some experiments, NK-specific *Kcnj8* KO and WT mice were injected intravenously with 100 µg poly I:C (Miltenyi Biotec) in 200 µL sterile PBS. After 24 h, splenocytes were harvested and NK cells were enriched with a negative NK cell isolation MACS kit, according to the manufacturer’s protocol. NK cells were co-incubated with RMA, RMA-S or RMA-RAE1y target cell lines in 1:1 ratio and treated with eBioscience Protein Transport Inhibitor Cocktail, according to manufacturer’s protocol. Cells were incubated for 48 h at 37 °C, and the frequency of CD107a expression in NK cells was determined with flow cytometry. In other experiments, after 4 days of tamoxifen injection, NK-specific *Kcnj8* KO and WT mice were injected intraperitoneally with 100 µg poly I:C (Miltenyi Biotec) in 200 µL sterile PBS. After 18h, splenocytes were harvested and NK cells were enriched with a negative NK cell isolation MACS kit, according to the manufacturer’s protocol. NK cells were co-incubated with YAC-1 target cells in 1:1 ratio with a test of conjugated CD107a antibody and treated with BD Biosciences GolgiStop Protein Transport Inhibitor Cocktail containing monensin, according to manufacturer’s protocol.

### *In vivo* killing assay with target cells

Three cell lines, RMA, RMA-S and RMA-RAE1y, were cultured in vitro in RPMI plus 10 % FBS at 37 °C. RMA cells were stained with Far Red Cell Proliferation kit (Thermo Fischer) while RMA-S and RMA-RAE1y cells were stained with CellTrace CFSE Cell Proliferation kit (ThermoFisher), according to manufacturer’s protocol. Equal amounts of each cell line were harvested and combined. The cells were administered intraperitoneally (in sterile PBS) to NK-specific *Kcnj8* KO mice and WT mice. The cells were harvested from the peritoneal cavity 48 h later, washed and quantified with flow cytometry by assessing CFSE, FarRed and h2-kb (to separate RMA-S and RMA-RAE1y).

### RNAscope in situ hybridization

Mouse spleens were carefully handled and fixed for 36-48h at room temperature with 4% paraformaldehyde (at pH 7.4 prepared in PBS) in a sufficiently large volume (at least 10x spleen volume) with gentle rocking ensuring that no air bubbles were in contact with the spleen. Tissue was processed by NYUMC’s Experimental Pathology Research Laboratory (RRID:SCR_017928), which included paraffin embedding, sectioning (3 µm thickness) and the fluorescence RNA in situ hybridization assays with RNAscope® technology using the RNAscope Multiplex Fluorescent V2 Assay (Advanced Cell Diagnostics, Newark, CA). Multispectral imaging was performed with a Akoya/PerkinElmer Vectra® instrument. The RNAscope™ LS 2.5 probes (Advanced Cell Diagnostics, Newark, CA) used included Mm-Kcnj8 (#411398), Mm-Abcc9 (#411378), Mm-Ncr1-C2 (#501729), Mm-Cd3e-C3 (#314728), and Mm-Adgre1-C4 (#460658). Opal dyes (Akoya Biosciences, MA) were used for secondary staining as follows: Opal 690 for C1 and Opal 570 for C3. DAPI was used for nuclear staining.

The spectrally unmixed fluorophores were demultiplexed using InForm software (Akoya/PerkinElmer). Cell segmentation, based on DAPI nuclear staining, and mRNA dots/cell were analyzed using HALO version 3.6 (Indica Labs, Albuquerque, NM). For nuclear detection the Nuclear Segmentation AI classifier was further trained with nuclei from the images used to quantify and labeled as ‘Nuclear_Segmentation_Mouse_spleen’. Each cell was defined as the nuclear segmentation plus 1 µm radius from the nuclear edge to account for cytoplasm. For detection of RNAscope dots the FISH module v3.2.3 was used with the parameter detailed in **Table S2**. Cells with one or more dots per probe were classified as positive for that probe.

### Patch clamp recordings

Mouse spleen NK cells were used for patch clamping within 8 hours after isolation. NK cells were seeded on a laminin-coated coverslip and allowed to attach for 5-10 min. Whole-cell patch clamping was performed by NYU Langone Health’s Ion Laboratory (RRID: SCR_021754) using an Axopatch-200B amplifier. Data were recorded with a Digidata 1550A and Clampex 10 software. Currents were low-pass filtered at 2 kHz with a 8 pole Bessel response, and data were acquired at 10 kHz (Digidata 1550A and Clampex 10.7; Molecular Devices). The bath solution consisted of (in mM) 140 NaCl, 5 KCl, 1 MgCl_2_, 1 CaCl_2_, 10 glucose, 10 HEPES, pH 7.4 adjusted with NaOH. Patch pipettes were made using borosilicate capillaries (1.5 mm O.D.; World Precision Instruments, Sarasota, Florida) and had resistances of ∼4 MΩ when filled with a solution consisting of (in mmol/L) 130 KCl, 5 EGTA, 1 CaCl_2_, 1 MgCl_2_, 10 HEPES, 0.1 Mg-ATP, 1 Na_2_-UDP, pH 7.2 adjusted with KOH. From a holding potential of −70 mV, 200 ms voltage steps were applied between −140 to 50 mV (10 mV increments) at 2 s intervals. Recordings were not corrected for the liquid junction potential (estimated to be 9.4 mV). Currents were calculated as the current density (pA/pF) by dividing the current by the cell capacitance. Data were analyzed using pClamp software and custom scripts written in Python 3.0.

### Measurement of intracellular Ca^2+^

Freshly isolated mouse splenic NK cells were plated at a density of 3.0-4.0 × 10^5^ cells/well in black, flat-bottom, 96-well plates and loaded with 1 μM Fluo4/AM (Life Technologies) for 30 min in complete RPMI medium. NK cells were pre-incubated for 15 min in Ca^2+^-free Dulbecco’s Phosphate Buffered Saline solution (DPBS. 1 mM MgCl2, 10 mM HEPES, pH 7.4) with pinacidil (100 µM), TRAM-34 (10 µM), PNU-37883A (10 µM) or with solvent only (<0.1% DMSO). Fluorescence was recorded at excitation/emission wavelengths of 485 nm/525 nm using a microplate reader equipped with a dispenser unit for adding compounds during the recording (FlexStation 3, Molecular Devices). Fluorescence intensities were recorded every 2 s for the duration of the experiment. Store depletion was induced by stimulating the cells with 1 μM thapsigargin in Ca^2+^-free DPBS solution, and Ca^2+^ influx was induced by adding an equal volume of 0.5 mM Ca^2+^ DPBS solution to the cells (for a final Ca^2+^ concentration of 200 µM). The fluorescence intensity of each well was normalized (F/F0) by dividing by the average baseline fluorescence intensity value prior to addition of thapsigargin (F0).

### Drugs used

We used pinacidil (Sigma Aldrich, St. Louis, MO), TRAM-34 Sigma Aldrich), PNU-37883A (Tocris Bioscience, Minneapolis, MN), glibenclamide (Sigma Aldrich), thapsigargin (Invitrogen), PMA/ Ionomycin (Biolegend), and Golgi Stop (Monensin) (BD Biosciences).

### Antibodies used

Antibodies used are shown in **Table S1**.

### Statistical analysis

When comparing two groups we used the Student’s t-test. For multiple group comparison we used 1-way ANOVA. If overall significance was achieved, ad-hoc pairwise comparison was performed using the Tukey t-test, or the Dunnett’s t-test when comparisons were made to a single control. Statistical significance was assumed at a p-value <0.05.

## Results

### *Kcnj8* is expressed in subsets of NK cells

Datasets from the Immunological Genome Project (ImmGen) mouse database show that *Kcnj8* is highly expressed in spleen CD27^-^CD11b^+^ NK cell subsets and, upon viral infection, is strongly upregulated in cytotoxic CD8^+^ T cells (**Figure 1A**). We confirmed the expression of *Kcnj8* mRNA by RT-PCR in both isolated mouse NK cells and in mouse brain (**Figure 1B**). Using RNAscope analysis, we found *Kcnj8* expression in 4.4% of cells in the mouse spleen (**Figure 1C and Figure S2**). Only a subset of *Ncr1*-postive NK cells (11.1%) had *Kcnj8* RNAscope dots. Thus, it appears that *Kcnj8* is not expressed in all of the NK cells in naïve mice.

**Figure 1:**
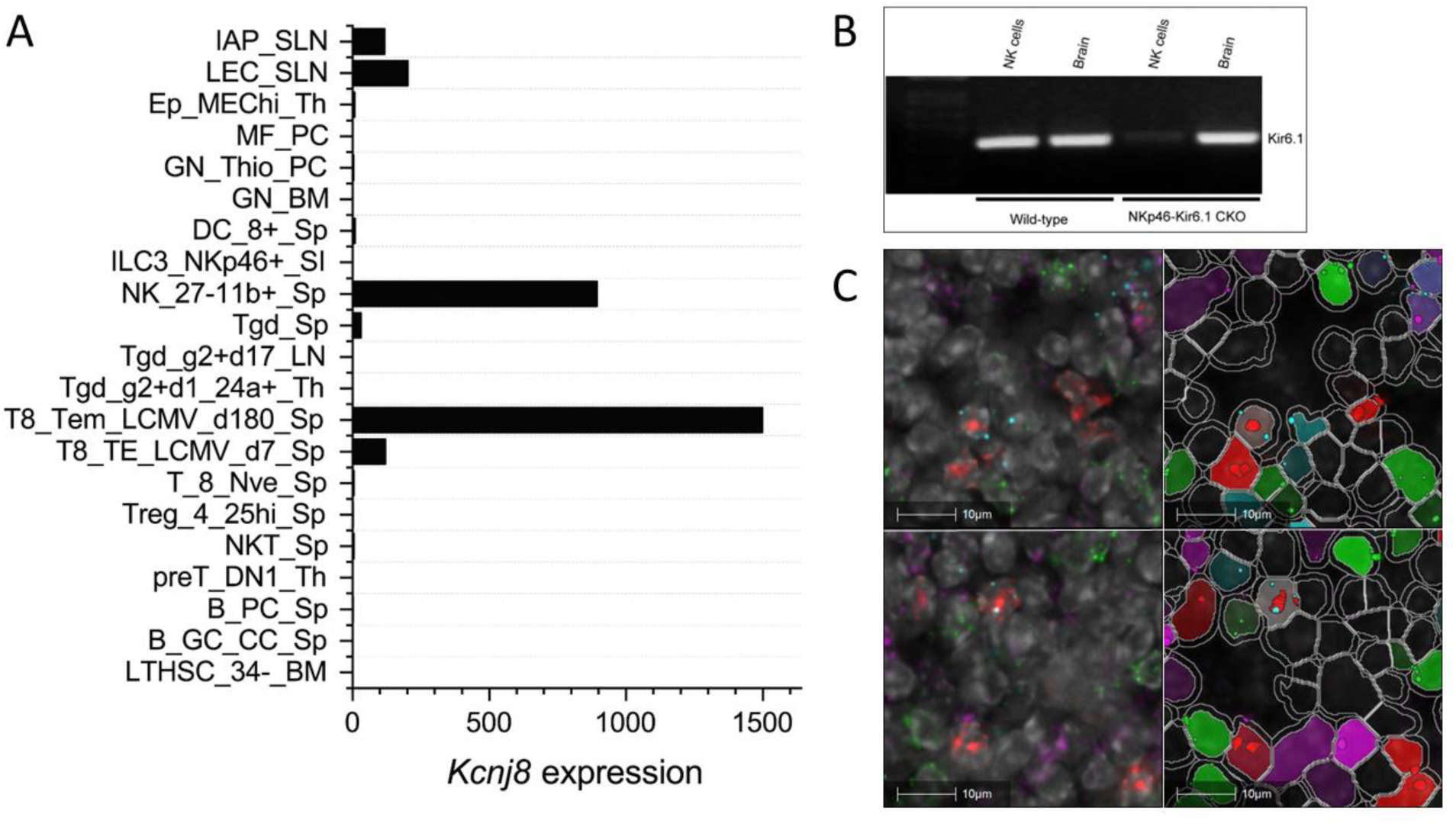
*Kcnj8* mRNA expression in NK cells. A) Kcnj8 is expressed in cytotoxic effector cells in mouse immune cells. Shown are ImmGen ULI RNA seq expression data of bone marrow hematopoietic stem cells (LTHSC_34-BM), splenic germinal center centrocytes (B_GC_CC_Sp), splenic plasma cells (B_PC_Sp), thymic preT DN1 cells (preT_DN1_Th), splenic natural killer T cells (NKT_Sp), splenic CD25hi TRegs (Treg_4_25hi), splenic naïve CD8+ T cells (Sp T_8_Nve_Sp), splenic CD8+ T cells 7 days after LCMV infection (T8_TE_LCMV_d7_Sp), splenic CD8+ T cells 180 days after LCMV infection (T8_Tem_LCMV_d180_Sp), immature thymocytes (Tgd_g2+d1_24a+_Th), Tgd_g2+d17_LN, total splenic dgT cells (Tgd_Sp), splenic CD27-/CD11b+ NK cells (NK_27-11b+_Sp), lamina propria NKp46+ ILC3 cells (ILC3_NKp46+_SI), splenic CD8+ cells (DC_8+_Sp), bone marrow neutorophils (GN_BM), thio-induced peritoneal neutrophils (GN_Thio_PC), peritoneal macrophages (MF_PC), thymic medullary epithelial cells (Ep_MEChi_Th), subcutaneous lymphatic endothelial cells (LEC_SLN), and subcutaneous lymphatic pericytes (IAP_SLN). Expression values are normalized by DESeq2. Not shown are immune cell types with expression levels below 0.1. Data are from http://rstats.immgen.org/Skyline/skyline.html. B) qRT-PCR data showing *Kcnj8* expression in isolated splenic NK cells and brain tissue from wild-type C57BL/6 mice and NK-cell specific NKp46-Kir6.1conditional knock-out mice. C) RNAscope data of a mouse spleen. RNAscope probes used were designed to localize NK cells (*Ncr1*; red), T cells (*Cd3*; green), macrophages (*Abgre1*; magenta), and cells expressing *Kcnj8*; light blue). Shown in the left are unprocessed images, with DAPI staining in gray scale. Cell segmentation was performed with DAPI as the cell marker and is shown on the right. Detected cells are outlined in white. Cells are pseudo colored based in expression of RNAscope dots. The top and bottom rows are two representative images.

### *Kcnj8* is expressed in mature and educated NK cells

To investigate *Kcnj8* expression in specific NK cell populations, we used a single cell RNA seq (scRNA seq) approach. NK cells were isolated from mouse spleens using a negative selection method, which largely excludes other cell types. Pooled cells from 3 mice were subjected to scRNA seq and analyzed using the Seurat package. A total of 15,878 cells were analyzed which yielded 16 individual cell clusters (**Figure 2A**). Clusters 0, 1, 2, and 3 (75.5 % of the total) were NK cells, based on the expression of NK marker genes *Klrb1c* (NK1.1), *Ncr1* (NKp46), *Cd27* (CD27) and *Itgam* (CD11b), and the lack of the T-cell marker gene *Cd3ε* (**Figure 2B**). The remaining ∼25 % of cells included minor populations of B cells, macrophages, NKT cells, CD8+ T cells, and other immune cells (**Table S3**).

**Figure 2:**
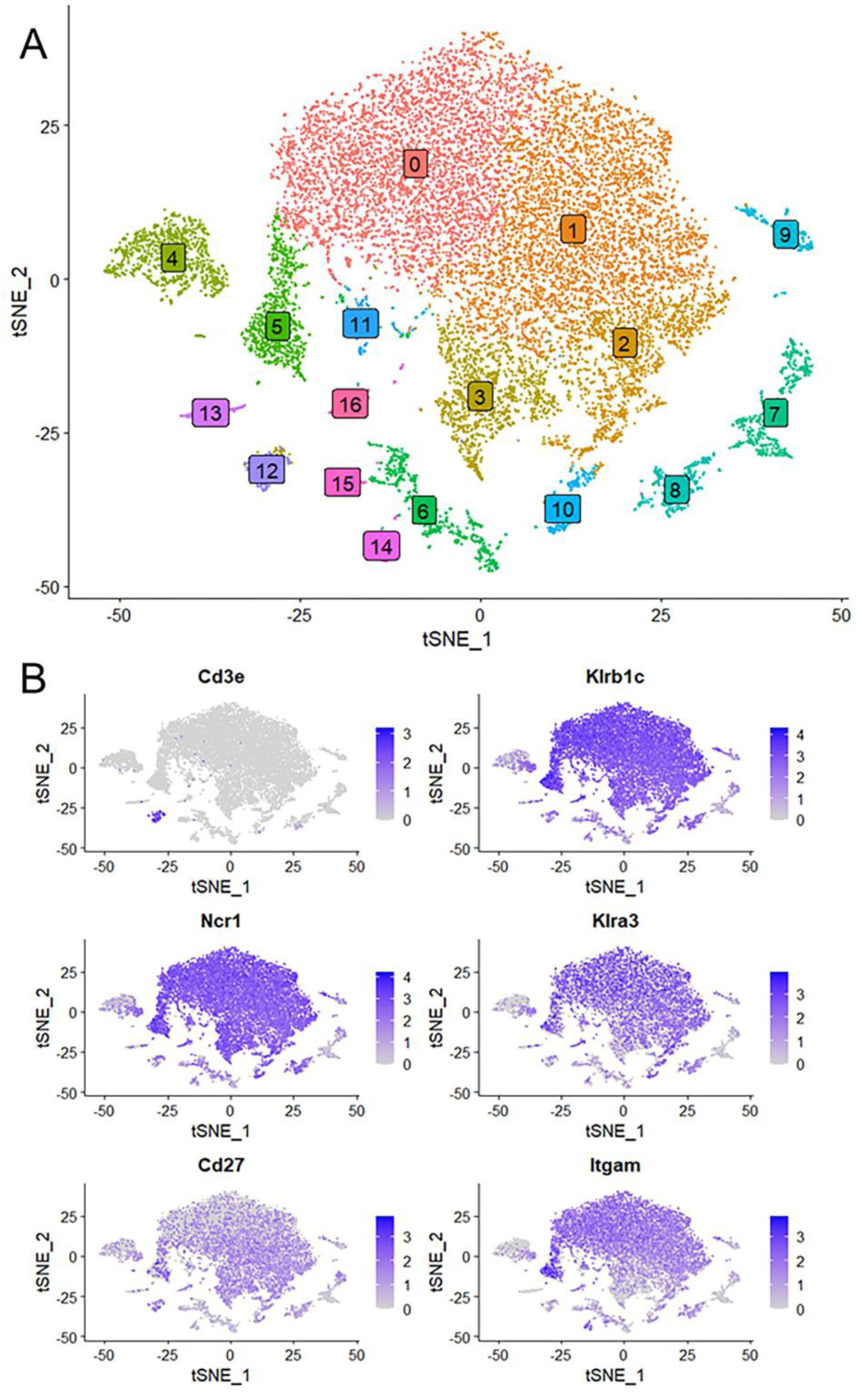
Single cell RNA seq data were obtained using NK cells of mouse spleen that were isolated using a negative selection method. Data analysis was performed with the Seurat package. A) Sixteen separate cell clusters were identified by graph-based clustering algorithm. Non-linear dimensional reduction was performed and data are displayed as t-distributed Stochastic Neighbor Embedding (tSNE) plots. B) Marker feature tSNE plots show lack of expression of *CD3e* (CD3E) in populations 0 to 3, and expression of the NK cell markers *Klrb1c* (NK1.1), *Ncr1* (NKp46), *Cd27* (CD27) and *Itgam* (CD11b). Also shown is high expression of *Klra3* (Ly49C) in mature NK cells (population 0).

We next focused on the major cell populations 0 to 3 that express the NK cell markers. As maturity progresses, NK cells transition through CD27^+^CD11b^-^, CD27^+^CD11b^+^ and CD27^-^ CD11b^+^ subsets, becoming increasingly become more cytotoxic during this progression (3). By the expression of these maturity markers, we identified populations 0 to 3 respectively to represent CD27^low^/CD11b^high^, CD27^high^/CD11b^high^, CD27^high^/CD11b^low^ and a minor population of CD27^low^/CD11b^low^ (**Figure 3A**). Thus, population 3 is the least mature, followed by populations 2, 1 and 0. Interestingly, *Kcnj8* tracked the expression of CD11b, demonstrating that its expression is highest in the mature NK cells (**Figure 3B**).

**Figure 3:**
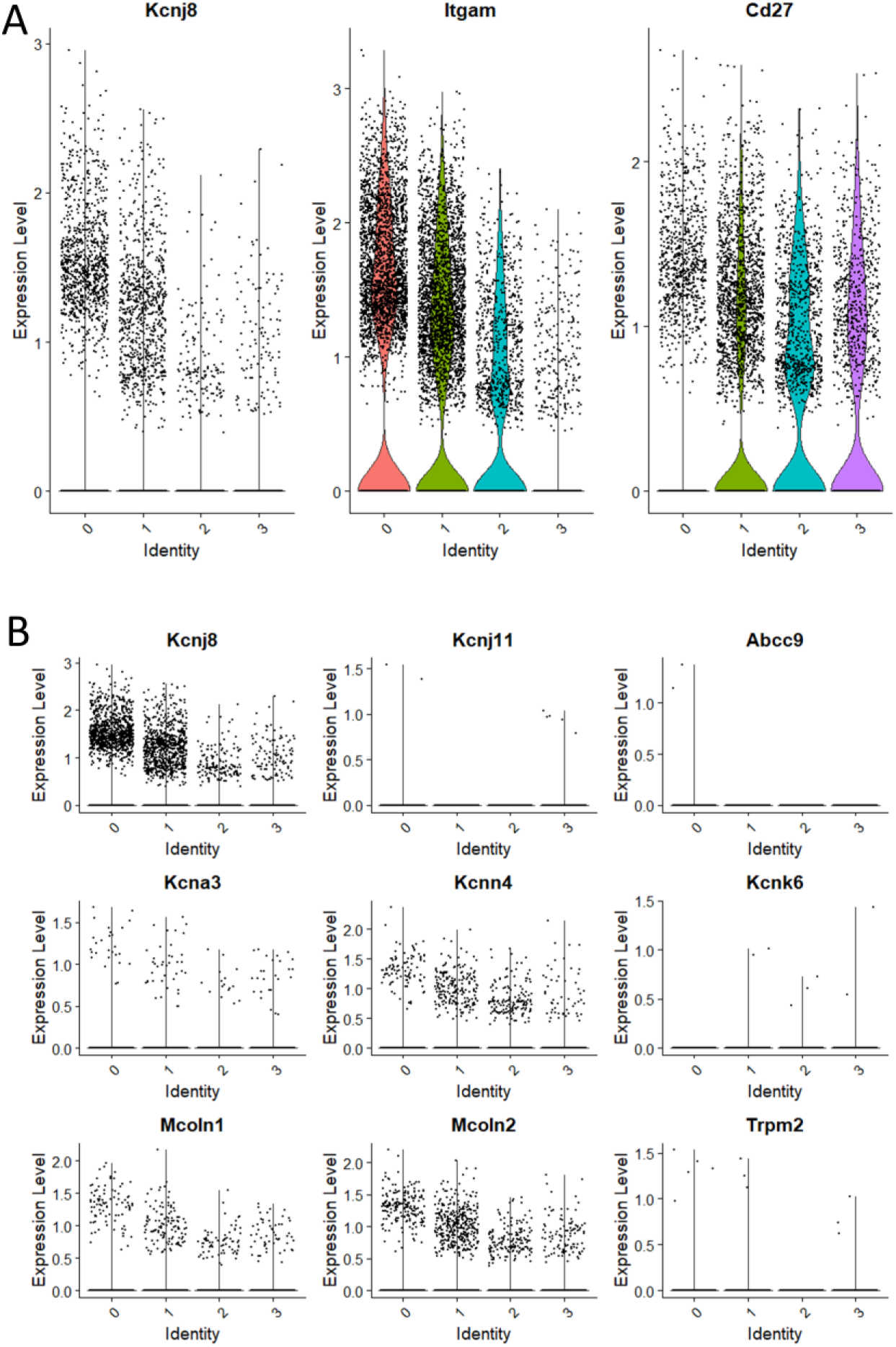
*Kcnj8* mRNA expression is highest in the mature CD11b+ NK cells in the mouse spleen. These data focus on populations 0 to 3 that are enriched in NK cell makers, identified in Figure 2. A) Single cell violin plots are shown for *Kcnj8* (Kir6.1), and the NK cell maturity markers *Itgam* (CD11b), and *Cd27* (CD27). B) Single cell violin plots of the scRNA seq data were generated to show the expression in populations 0 to 3 of ion channel genes *Kcnj8* (Kir6.1), *Kcnj11* (Kir6.2), *Abcc9* (SUR2), *Kcna3* (Kv1.3), *Kcnn4* (Kv1.3), *Kcnk6* (TWIK-2), *Mcoln1* (TRPML1), *Mcoln2* (TRPML2), and *Trpm2* (TRPM2). Note that *Abcc8* (SUR1) had no gene counts in this dataset.

We compared the expression of *Kcnj8* to other ion channels that have previously been identified in NK cells (**Figure 3B**). The expression of *Kcnj8* (Kir6.1) far exceeds those of K^+^ channels, including *Kcnj11* (Kir6.2), *Abcc9* (SUR2), *Kcna3* (Kv1.3), *Kcnn4* (Kv1.3), *Kcnk6* (TWIK-2), *Mcoln1* (TRPML1), *Mcoln2* (TRPML2), and *Trpm2* (TRPM2). *Abcc8* (SUR1) was not present in this NK cell dataset. High expression of *Kcnj8* in CD27^low^CD11b^high^ splenic NK cells can also observed by reexamining data in public databases (**Figure S3**) and other independent reports observed high expression of *Kcnj8* in mature mouse NK cell subsets (33–35), including educated NK cells (36, 37).

Uterine NK cells follow a different developmental trajectory from that of peripheral blood NK (pbNK) cells in both humans and mice. To analyze transcriptional changes downstream of educated uNK cells, we purified NKG2A^+^ and Ly49C/I^+^ uNK cells and analyzed differential gene expression (DEG) by comparing their transcriptomes with that of purified, uneducated NKG2A^-^Ly49C/I^-^ uNK cells. The data show that *Kcnj8* is expressed at elevated levels in NKG2A^+^ or Ly49C/I^+^ educated uNK cells, alongside known markers of NK-cell education *Cd266* (DNAM-1). Other highly expressed DEG include cytokines and chemokines (e.g: Ccl5, Il15), effector molecules (e.g. *Ifng*, *Xcl1, Gzm*), transcription factors, and signaling molecules (**Figure 4**). Overall, consistent with the data obtained with splenic NK cells, educated uNK cells expressed elevated levels of *Kcnj8*.

**Figure 4:**
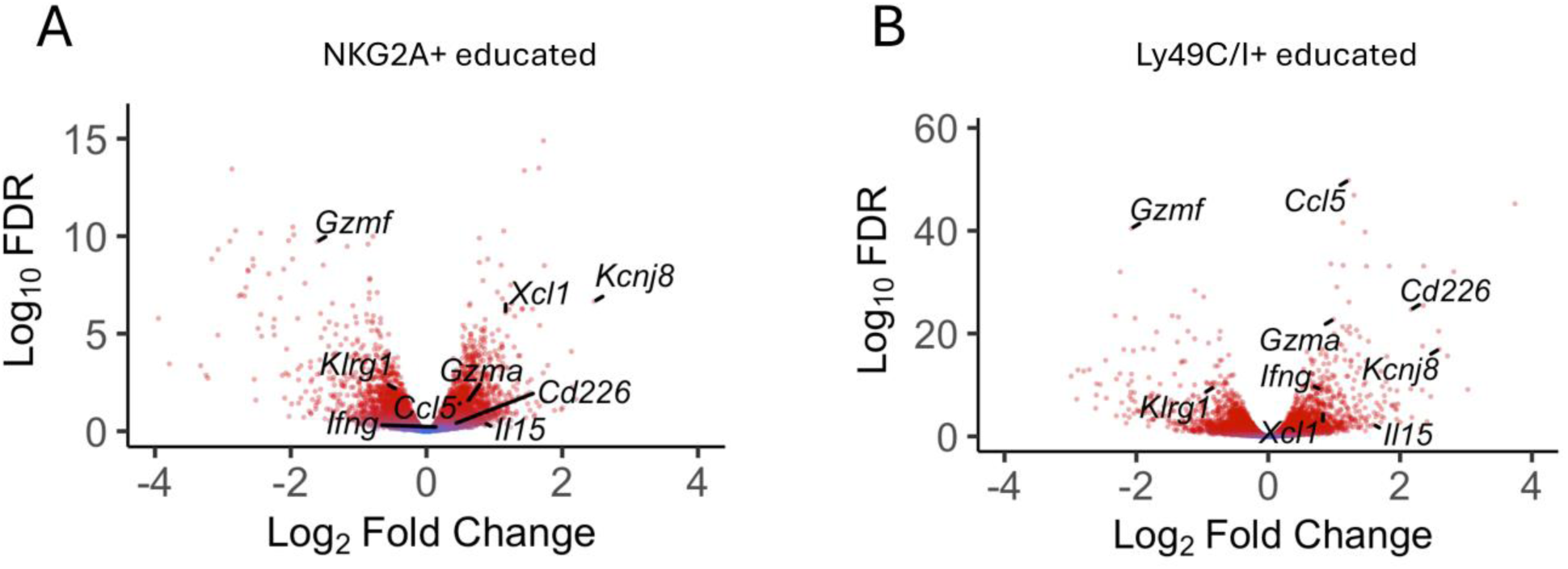
*Kcnj8* is expressed at higher levels in mature, educated uterine NK cells. A) Volcano plot showing selected DEGs in NKG2A+ (n=9) and LY49C/I+ (n=6) educated uNK compared to uneducated uNK cells (LY49C/I-NKG2A- n=9). Red color indicates statistical significance (FDR<0.05). Fold change expressed in log2 relative to uneducated uNK cells.

### K_ATP_ channel recordings in mouse splenic NK cells

We next performed patch clamp experiments to determine whether K_ATP_ channel currents can be measured in NK cells. NK cells were isolated from mouse spleen using a negative selection method and subjected to whole-cell patch clamping on the same day. Typical whole-cell patch clamp recordings are shown in **Figure 5**. Two types of current profiles were present in these cells. In 6 of 10 cells, there was a time-dependent activation of currents during depolarization and the currents were strongly outward rectifying (**Figure 5A & B**). The time-dependent profile is indicative of the presence of a voltage-gated K^+^ current. We used PNU-37883A in these experiments since it is a Kir6.1 pore blocker (36, 37). Neither the K_ATP_ channel opener pinacidil (100 µM) nor PNU-37883A (10 µM) had significant effects on the recorded currents. Four of the 10 cells, by contrast, exhibited little time dependence and the currents were relatively linear as a function of voltage (**Figure 5C & D**). Pinacidil had no appreciable effects on these currents, but PNU-37883A blocked both inward and outward currents. At −100 mV, for example, PNU-37883A decreased the current by −46 ± 6.7 % (n=4; p<0.01; paired-t-test). These data are consistent with the presence of a Kir6.1-containing K_ATP_ channel in a subset of NK cells that is sensitive to the channel blocker PNU-37883A.

**Figure 5:**
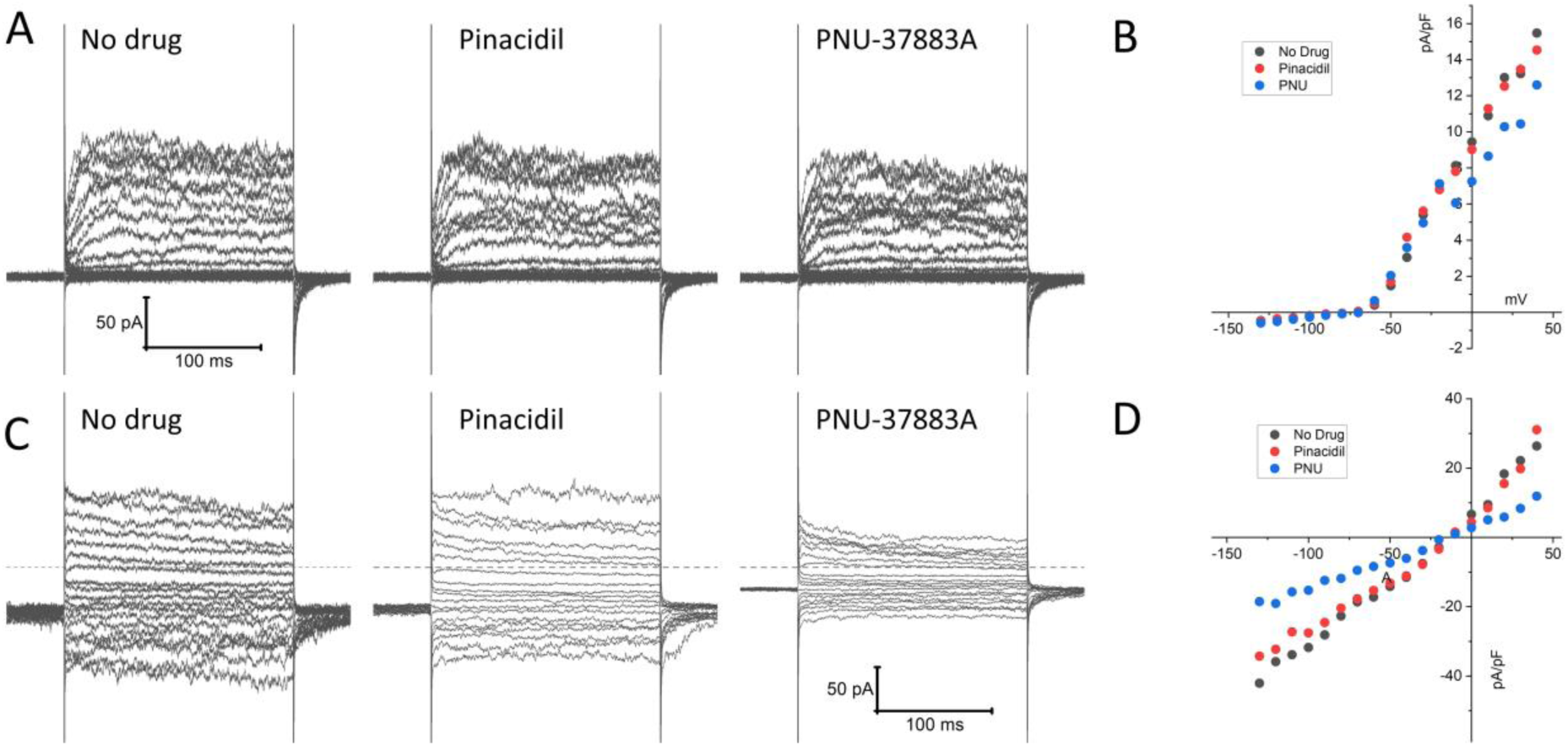
Patch clamp current recordings in isolated mouse spleen NK cells. NK cells, isolated from wild-type mouse splenocytes by negative selection, were subjected to whole-cell patch clamp recordings. A) An example of recordings of a cell that exhibited time-dependent currents upon depolarization with little inward rectification. Shown on the left are recordings made in the absence of drug, 3 min after application of 100 µM pinacidil, and 3 min after application of 10 µM PNU-37883A (in the continued presence of pinacidil). B) Depicted is the current, measured at the end of the voltage step, as a function of applied voltage of this cell, demonstrating pronounced outward rectification. C) A typical example of a cell with a distinctly different current profile, that showed little time dependence and the presence of strong inward currents upon hyperpolarization. D) The current-voltage relationship of this cell was essentially linear, and the current was strongly blocked by PNU-37883A.

### *Kcnj8* does not regulate degranulation of NK cells

We first assessed the effects of pharmacological K_ATP_ channels intervention on NK cell degranulation. Surface expression of CD107a (LAMP-1) was measured by flow cytometry as an index of degranulation. WT mice were compared with tamoxifen-inducible NK cell-specific *Kcnj8* deficiency. Mice were pretreated with poly I:C, a synthetic dsRNA that strongly stimulates innate immunity. NK cells were isolated from the spleens and were co-incubated for 2 h with YAC-1 target cells in the presence of the K_ATP_ channel opener (30 µM pinacidil), a K_ATP_ channel blocker (1 µM glibenclamide). As a positive control, cells were incubated with a combination of phorbol 12-myristate 13-acetate (PMA; 80 nM) and ionomycin (1.3 µM), which is known to stimulate NK cell degranulation (38). A negative control consisted of the drug solvent only (<0.01 % DMSO). Neither the K_ATP_ channel opener nor the blocker affected degranulation in this *in vitro* assay (**Figure 6A and B**). Moreover, there was no significant difference when comparing the WT and KO groups (2W ANOVA; p = 0.794).

**Figure 6:**
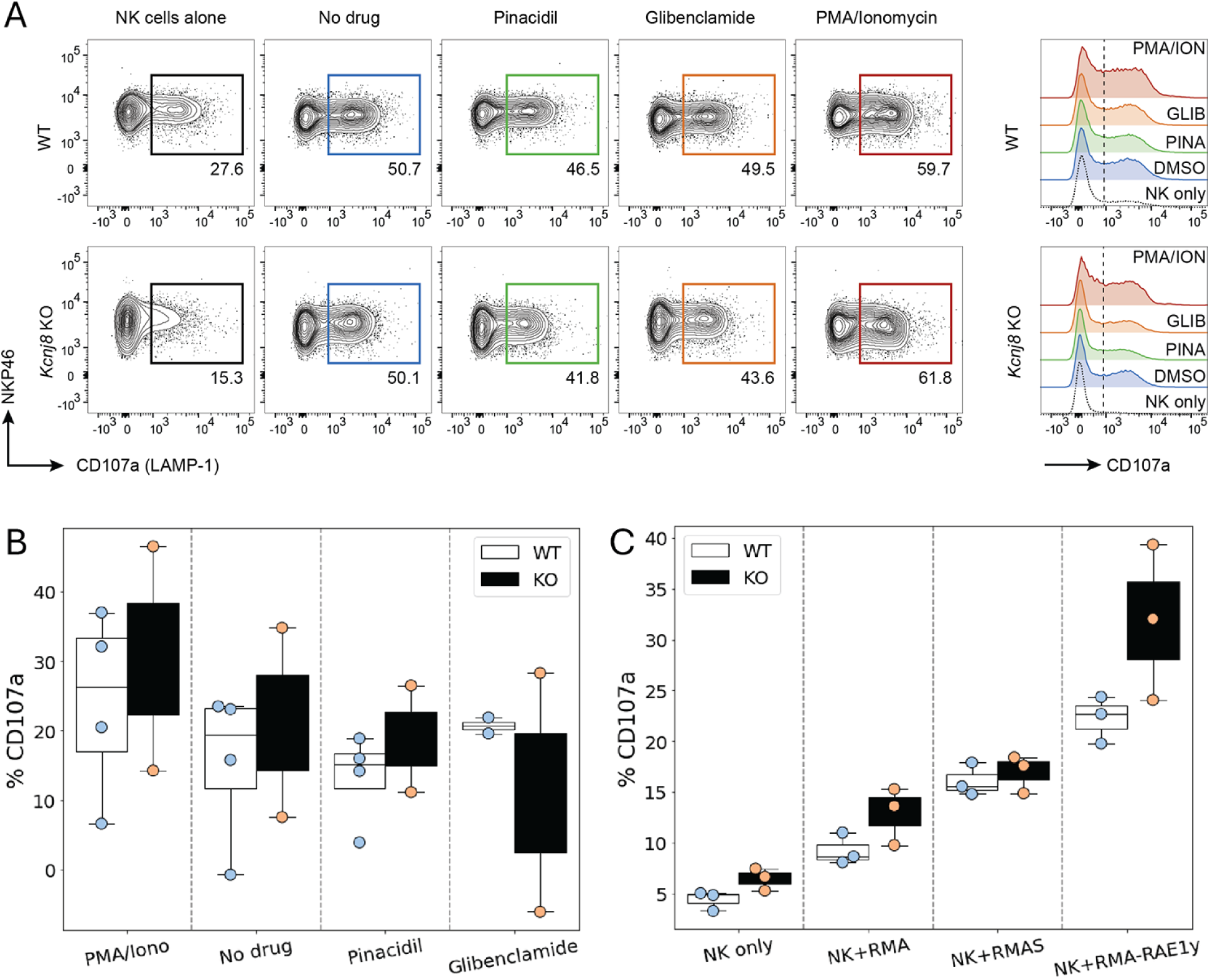
NK cell degranulate effectively in both WT and NK-cell specific *Kcnj8* KO mice. A) Flow cytometry gating strategy to detect CD107a+ (degranulating) NK cells in a co-culture of NK cells and YAC-1 cells. NK cells, isolated from mouse spleen, were co-cultured for 2 h with YAC-1 target cells in the presence of the K_ATP_ channel opener (30 µM pinacidil), a K_ATP_ channel blocker (1 µM glibenclamide). Phorbol 12-myristate 13-acetate (PMA; 80 nM) and ionomycin (1.3 µM) was used as a positive control. The no drug group contained solvent only (<0.01 % DMSO). B) Summary data of all experiments using the K_ATP_ channel drugs. C) Spleen NK cells isolated from WT or NK-specific *Kcnj8* KO mice were co-cultured with or without the target cell lines, RMA, RMA-S and RMA-RAE1y. Flow cytometry was performed to detect the percentage of CD107a+ NK cells (as a surrogate marker of degranulation). The data are from 3 separate experiments.

We next assessed the impact of *Kcnj8* on NK cell degranulation against different types of target cells. We used three types of lymphoma cells: parental RMA cells expressing MHC class I molecules; mutant RMA-S cells expressing very low MHC class I molecules and mutant RMA-Rae1γ cells expressing MHC class I molecules along with the NKG2D ligand. The Rae1γ. RMA-S cells and RMA-Rae1γ cells elicit stronger NK cell activation than the RMA cells (39, 40). We compared wild-type mice with mice constitutively deficient of *Kcnj8* in NK cells. In both groups, NK cells were pre-stimulated by injecting poly(I:C). After 2 days, NK-enriched splenocytes were co-incubated with either RMA, RMA-S or RMA-Rae1γ cells. The was quantified after 2 hours by flow cytometry (n=3-4, repeated twice). As expected, NK when compared to RMA cells (**Figure 6C**). There was, however, no statistical difference when comparing the genotypes (2W ANOVA; p=0.860).

Taken together, our data show that NK cells can degranulate effectively in the absence of *Kcnj8*. It is possible, however, that subsets of NK cells may be differentially affected,

### *Kcnj8* is not required for tumor rejection by NK cells

In a next experiment, we tested the efficiency of NK cells to reject tumor cells *in vivo*. We administered an equal mix of RMA, RMA-S and RMA-Rae1γ cells intraperitonially to control mice and mice constitutively deficient of *Kcnj8* in NK cells. After 48 hours, tumor cells were harvested and quantified. Very few MHC-deficient RMA-S cells were present in both groups, suggesting they had been rejected by NK cells upon missing-self recognition (n=3-4, repeated twice). We focused therefore on the ratio of RMA to RMA-Rae1γ cells. By flow cytometry, we detected several different cell populations in the peritoneal cavity (**Figure 7A**). The RMA and RMA-Rae1γ cells tumor cells were recovered in approximately the same ratio in the control and KO mice, suggesting no obvious defect in recognition of induced self in the KO mice (**Figure 7B**). These results suggest that the *in vivo* anti-tumor NK-cell response is effective in KO mice. Therefore, *Kcnj8* deficiency does not seem to affect overall NK-cell cytotoxicity under these specific conditions or tumor cells tested.

**Figure 7:**
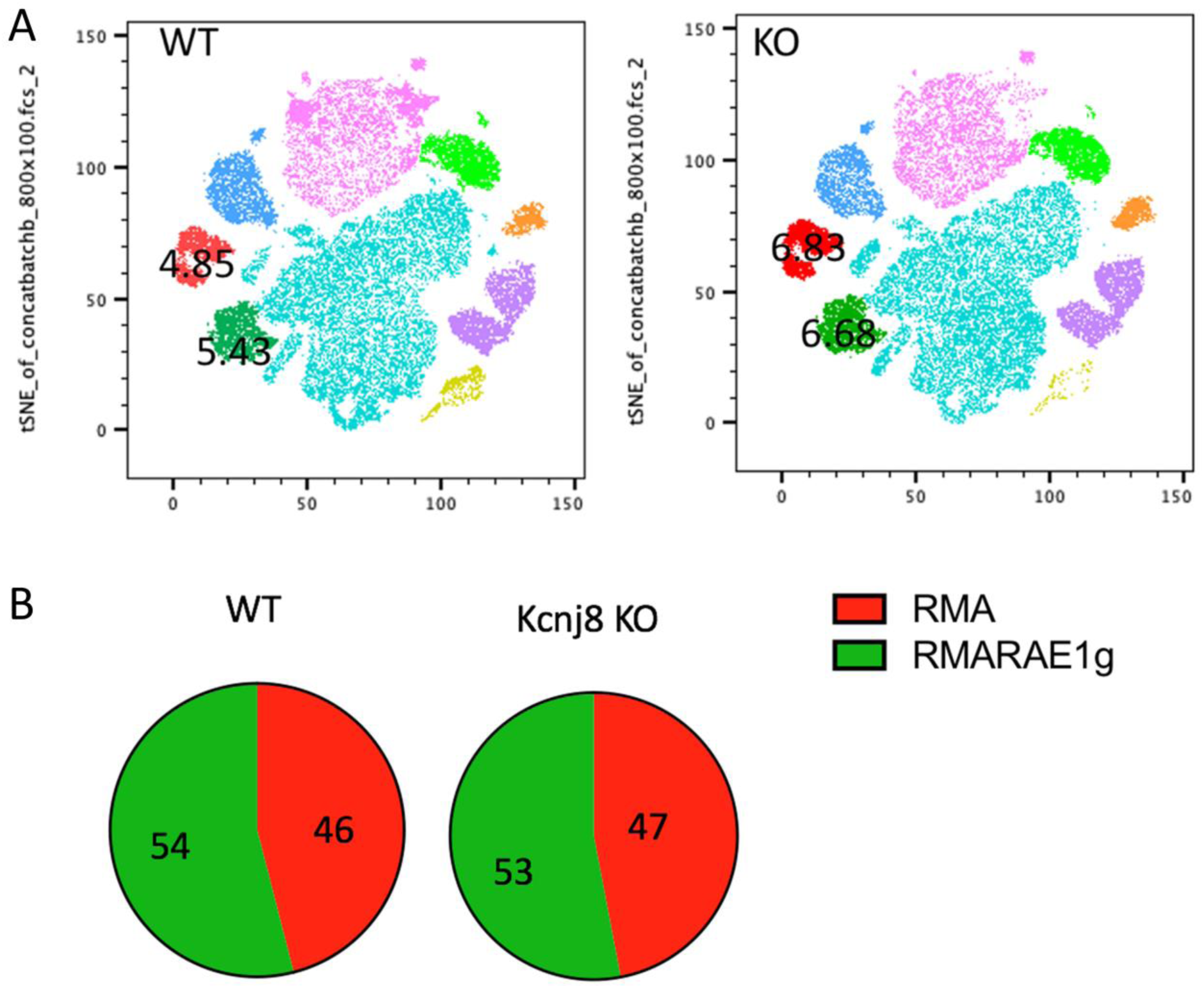
NK cells reject tumor cells equally *in vivo* in WT and NK-specified *Kcnj8* KO mice. Mice were injected I.P. with a mixture of RMA-S, RMA and RMA-RAE1y cells, pre-stained with fluorescent markers for later visualization and identification. The RMA-Rae1γ cells are preferential NK cell targets due to high expression of MHC class I molecules. After 48 h, cells were isolated from the peritoneum and subjected to flow cytometry to quantify the relative amounts of target cells. A) Cells from the peritoneal fluid of four WT and four NK-specified *Kcnj8* KO mice were downsampled to equal cell counts of viable cells per group (40.000 cells per group) in flowjo and concatenated to a single file prior to unbiased dimensionality reduction approach, visualised as tSNE plots. The clusters were determined by protein expression levels of forward and side scatter (FSC & SSC), H2Kb, CD3, CFSE, FarRed, NK1.1 and CD69. RMA-S cells were not detected. The RMA (red) and RMA-RAE1y (green) cell populations each comprised about 5% of the number of cells in both the WT and KO mice. B) Depicted is the percentage of RMA and RMA-RAE1y cells found in peritoneal fluid of WT and KO mice. Experiment was repeated once.

### No role for K_ATP_ channels in store-operated Ca^2+^ entry in mouse NK cells

Degranulation is dependent on store-operated Ca^2+^ entry (SOCE) into human NK cells (21), The Ca^2+^-activated K^+^ channel KCa3.1 regulates SOCE by preventing SOCE-induced membrane depolarization, thereby maintaining Ca^2+^ influx (16). Given the high expression of the *Kcnj8* in mouse cytotoxic NK cells, we tested whether K_ATP_ channels may have a similar role in SOCE in mouse NK cells. We used a well-described protocol to deplete Ca^2+^ from the endoplasmic reticulum (ER) with the sarco-endoplasmic reticulum Ca^2+^ ATPase (SERCA) inhibitor thapsigargin (TG), which triggers SOCE through Ca^2+^ release-activated Ca^2+^ (CRAC) channels (41). We recorded SOCE by measuring cytosolic Ca^2+^ with Fluo-4/AM while depleting the ER stores, and following the addition of 200 µM extracellular Ca^2+^ to initiate SOCE (**Figure 8A**). The magnitude of the SOCE response was unaffected by the K_ATP_ channel opener pinacidil or the Kir6.1-blocker PNU-37883A (**Figure 8A & B**). Blocking KCa3.1 with TRAM-34 also had no effect on SOCE, which is consistent with the low *Kcnn4* (KCa3.1) mRNA counts in the mouse spleen NK cells (**Figure 3C and Figure S3**). These data suggest that neither KCa3.1 nor K_ATP_ channels contribute to SOCE in mouse splenic NK cells.

**Figure 8:**
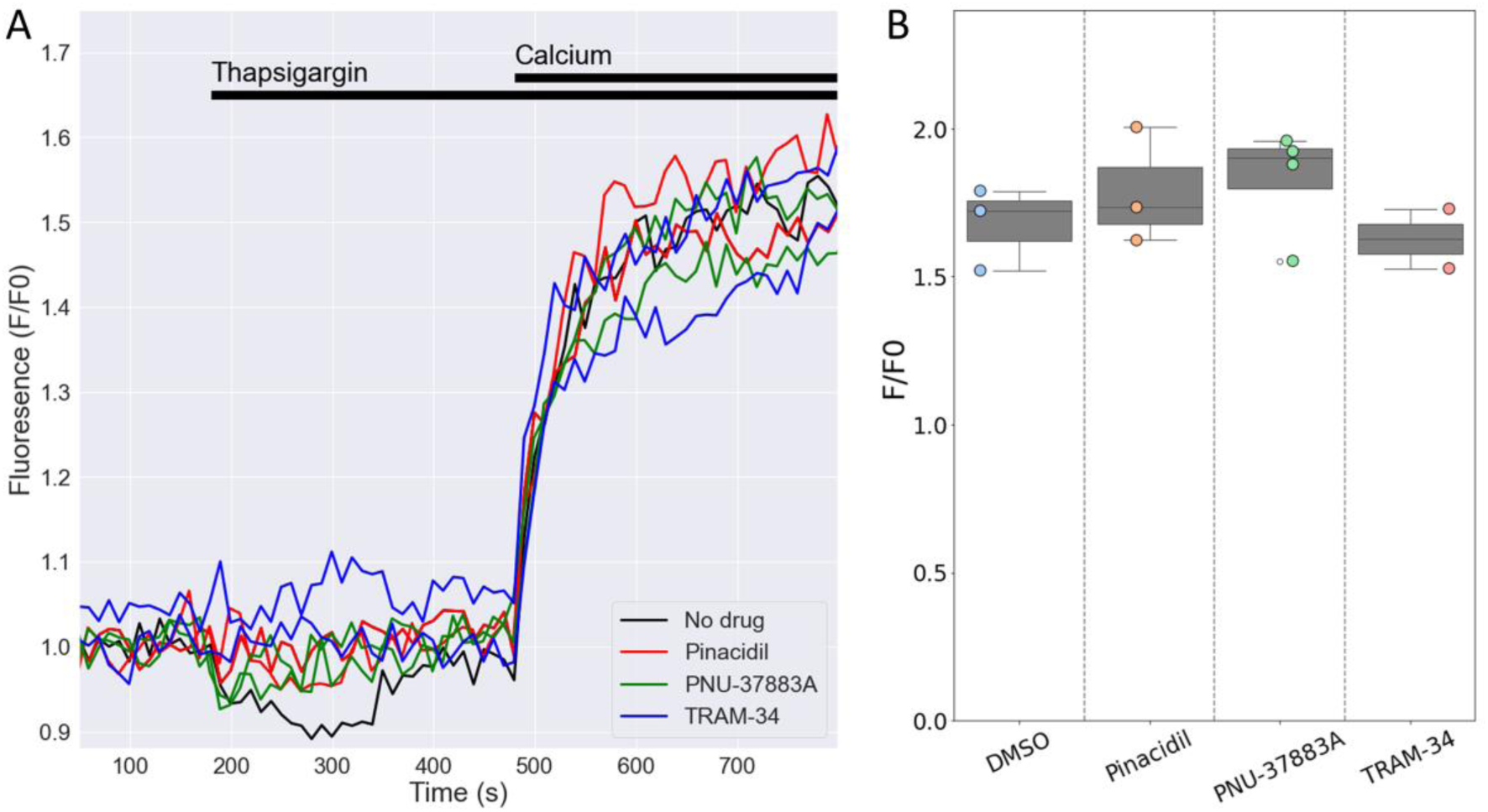
Openers or blockers of K_ATP_ channels do not affect store-operated Ca^2+^ entry (SOCE) in mouse NK cells. Cytosolic Ca^2+^ of isolated splenic NK cells was recorded with Fluo4/AM. A) Depicted are representative F/F0 traces in different wells of the same experiment with NK cells pretreated for 15 min with pinacidil (100 µM), TRAM-34 (10 µM), PNU-37883A (10 µM) or with no drug (<0.1% DMSO). Store depletion was accomplished by 1 μM thapsigargin and SOCE was initiated by adding 200 µM CaCl_2_ to the external solution. B) Summary data of experiments performed on different days (n=3 mice). p=0.84 with 1W-ANOVA.

### Effects of *Kcnj8* deficiency on the transcriptome of NK cells

In order to better understand the consequences of NK cell *Kcnj8* deficiency, we performed a bulk RNA seq experiment of freshly isolated splenocytes, with NK cells isolated by cell sorting using CD27 and CD11b antibodies. RNA seq experiments were performed with sorted NK cells from wild-type mice and tamoxifen-inducible NK cell-specific *Kcnj8* deficiency (n=2 each). The expectation was that these data may point to underlying transcriptional changes and pathways affected by the absence of *Kcnj8*. The read counts were variable in the immature CD27-/CD11b- population due to low cell yields and these data were therefore excluded from analysis. A principal component analysis of the remaining three populations (**Figure 9A**) demonstrated that NK cell populations CD27+/CD11b-, CD27+/CD11b+ and CD27-/CD11b+ (respectively populations 2, 1 and 0 in order of maturity) segregated separately, indicative of transcriptional differences between these cell types. The variance between genotypes were small, demonstrating that *Kcnj8* deficiency led to specific transcriptional changes. When mapping the individual reads to *Kcnj8* on the mouse genome, it is clear that reads within exon 2 are largely absent in the *Kcnj8* knockout mice, as expected from the knockout strategy (**Figure 9B**). The normalized reads for *Cd27* (CD27) and *Itgam* (CD11b), which verifies the nature of the NK cell populations are shown in **Figure 9C**. Also shown are the normalized reads of *Kcnj8* in the different groups, which makes it evident that the mature CD27-CD11b+ NK cells express significantly elevated levels of *Kcnj8* compared to the less mature CD27+/CD11b+ and CD27+/CD11b- populations in WT mice. Given that *Kcnj8* expression is so much elevated in the CD27-/CD11b+ population, we performed a differential gene expression analysis in this population by comparing the WT and KO genotypes. A total of 29 genes were differentially expressed, with 16 upregulated and 13 downregulated (**Table 1**). A gene ontology analysis reveals affected biological processes to be involved that include phospholipase C activity, cytokine production and JAK-STAT signaling (**Figure S4**).

**Figure 9:**
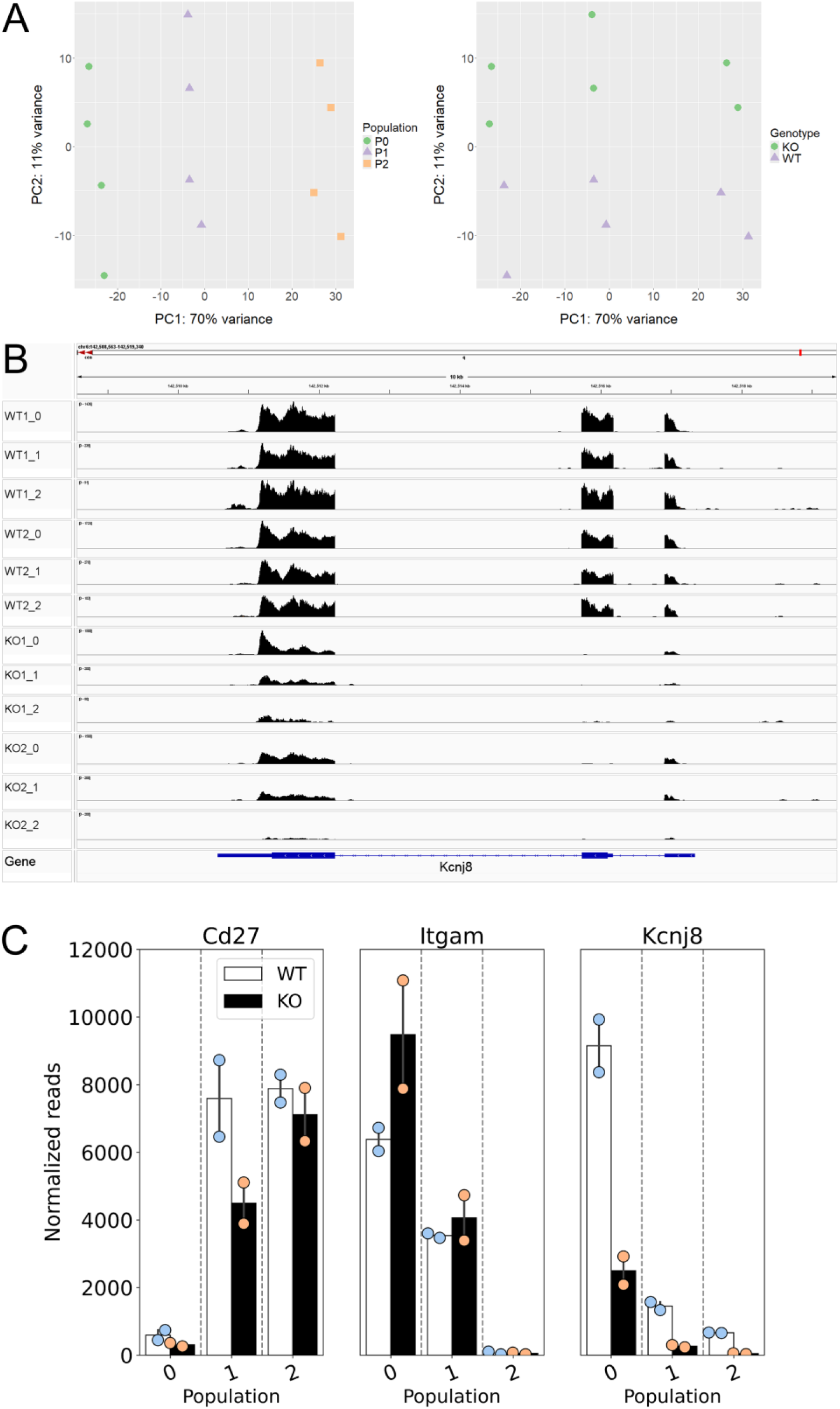
Data of a bulk RNA seq experiment performed with mouse splenic cells, NK cells isolated by cell sorting by first using NK markers NK1.1 and NKp46, then NK cell maturity markers CD27 and CD11b (n=2 each of WT and tamoxifen-induced NK cell-specific *Kcnj8* KO mice). Normalized reads and differentially expressed genes were calculated using the DEseq2 package. Normalized gene expression was averaged within populations. A) Principal component analysis demonstrates segregation of the three populations 2, 1 and 0 (respectively CD27+/CD11b-, CD27+/CD11b+ and CD27-/CD11b+). B) Mapping of RNA seq reads against the mouse mm10 genome was performed with IGV (Integrative Genomics Viewer). Note that the *Kcnj8* gene at the bottom of the panel is oriented in the reverse direction (3’ to 5’). C) Normalized *Cd27* (CD27), *Itgam* (CD11b), and *Kcnj8* (Kir6.1) reads in the three different populations for wild-type (WT) and mice with NK cell-specific *Kcnj8* deficiency (p<0.05 with the Mann-Whitney Rank Sum Test). Note that, although *Kcnj8* reads remain in the KO mice, they are from exons 1 and 3 as shown in panel B.

**Table 1:** Differentially expressed genes in the CD27^-^/CD11b^+^ population of NK cells.

| Gene | Name | baseMean | log2FoldChange | lfcSE | pvalue | padj |
| --- | --- | --- | --- | --- | --- | --- |
| *Uty* | ubiquitously transcribed tetratricopeptide repeat containing, Y-linked | 224.1 | -9.54 | 1.38 | 9.47E-12 | 1.59E-08 |
| *Kdm5d* | lysine demethylase 5D | 296.1 | -9.12 | 1.04 | 1.87E-17 | 4.05E-14 |
| *Capn11* | calpain 11 | 33.5 | -8.60 | 2.94 | 3.37E-07 | 0.000243 |
| *Ddx3y* | DEAD box helicase 3, Y-linked | 1034.6 | -8.14 | 0.53 | 6.78E-54 | 1.03E-49 |
| *Ighg2c* | immunoglobulin heavy constant gamma 2C | 747.5 | -6.69 | 0.51 | 4.14E-40 | 2.09E-36 |
| *Slpi* | secretory leukocyte peptidase inhibitor | 54.4 | -4.31 | 0.98 | 4.92E-07 | 0.00031 |
| *Kit* | KIT proto-oncogene receptor tyrosine kinase | 946.6 | -3.55 | 0.41 | 8.97E-20 | 2.26E-16 |
| *Dtx1* | deltex 1, E3 ubiquitin ligase | 328.1 | -2.91 | 0.43 | 3.01E-13 | 5.69E-10 |
| *H2-Ab1* | histocompatibility 2, class II antigen A, beta 1 | 346.1 | -2.89 | 0.56 | 1.05E-08 | 1.06E-05 |
| *Cd74* | CD74 antigen (invariant polypeptide of major histocompatibility complex, class II antigen-associated) | 547.9 | -2.86 | 0.48 | 1.22E-10 | 1.84E-07 |
| *Ltb* | lymphotoxin B | 198.1 | -2.51 | 0.53 | 7.65E-08 | 6.44E-05 |
| *Hk3* | hexokinase 3 | 201.0 | -2.28 | 0.47 | 5.25E-08 | 4.96E-05 |
| *Il7r* | interleukin 7 receptor | 216.8 | -2.21 | 0.51 | 4.87E-07 | 0.00031 |
| *Adgrg5* | adhesion G protein-coupled receptor G5 | 564.4 | -1.83 | 0.38 | 5.91E-08 | 5.26E-05 |
| *Kcnj8* | potassium inwardly-rectifying channel, subfamily J, member 8 | 4795.2 | -1.71 | 0.32 | 2.59E-09 | 3.02E-06 |
| *Tcf7* | transcription factor 7, T cell specific | 1320.5 | -1.59 | 0.36 | 3.13E-07 | 0.000237 |
| *Rhpn1* | rhophilin, Rho GTPase binding protein 1 | 2733.8 | 1.03 | 0.36 | 0.000125 | 0.045062 |
| *Tigit* | T cell immunoreceptor with Ig and ITIM domains | 707.1 | 1.31 | 0.37 | 1.58E-05 | 0.007478 |
| *Fam234b* | family with sequence similarity 234, member B | 434.6 | 1.55 | 0.38 | 1.60E-06 | 0.000837 |
| *Serpinb9b* | serine (or cysteine) peptidase inhibitor, clade B, member 9b | 10862.6 | 1.58 | 0.29 | 2.57E-09 | 3.02E-06 |
| *Frmd5* | FERM domain containing 5 | 56.8 | 2.48 | 0.80 | 6.94E-05 | 0.028408 |
| *Gp6* | glycoprotein 6 platelet | 81.3 | 2.94 | 0.75 | 3.72E-06 | 0.001878 |
| *Rab4a* | RAB4A, member RAS oncogene family | 76.8 | 3.30 | 0.73 | 2.76E-07 | 0.00022 |
| *Esr1* | estrogen receptor 1 (alpha) | 829.5 | 3.32 | 0.38 | 5.52E-20 | 1.67E-16 |
| *Emp1* | epithelial membrane protein 1 | 1643.9 | 4.80 | 0.44 | 1.78E-29 | 6.73E-26 |
| *Firre* | functional intergenic repeating RNA element | 62.9 | 4.95 | 0.92 | 6.44E-09 | 6.96E-06 |
| *Sh2d1b2* | SH2 domain containing 1B2 | 30.0 | 5.75 | 1.76 | 0.000147 | 0.049454 |
| *H2af-ps2* |  | 31.6 | 6.02 | 1.61 | 4.89E-05 | 0.021788 |
| *Xist* | inactive X specific transcripts | 825.3 | 8.16 | 0.60 | 1.10E-42 | 8.34E-39 |

Bulk RNA seq data are from mouse splenocytes, sorted by expression of CD27 and CD11b. DEseq2 analysis was performed to compare wild-type with Kir6.1 KO mice (n=2 each). Shown here are differentially expressed genes, defined by a log2fold value larger than 1.0 or smaller than -1.0; with an adjusted p-value < 0.05.

### *Kcnj8*-deficient NK cells fail to reach full maturity

Among the downregulated genes in the CD27-/CD11b+ NK cells in the *Kcnj8* KO were IL7R, Lymphotoxin B, and KIT proto-oncogene receptor tyrosine kinase (c-Kit). These all have key roles in NK cell development (42–44). We therefore investigated the possibility that *Kcnj8* may be involved with NK cell maturity, marked by the expression of surface markers CD27 and CD11b in bone marrow and spleen (3). We compared wild-type mice with mice with NK-cell specific *Kcnj8* ablation. First, we stained bone marrow NKp46^+^NK1.1^+^ NK cells of control and constitutive NK-specific *Kcnj8*-deficient mice (KO, n=3 each) with CD27, CD11b, DNAM-1, KLRG1 and CD49b to define discrete developmental stages in both groups. The subset (**Figure 10A**), and a reduction in mature KLRG1+ NK cells (**Figure S5**), whereas other populations were unchanged. We repeated this experiment with isolated splenic NK cells using wild-type and tamoxifen-induced NK cell-specific *Kcnj8* deficiency (**Figure 10 B&C**). As expected, the number of double negative (DN; CD27^−^CD11b^−^) immature NK cells was very low in the spleen. The major NK cell populations (in order of maturity) of wild-type mice were CD27^+^CD11b^−^, CD27^+^CD11b^+^ and CD27^-^CD11b^+^ NK cells. The total number of splenic NK cells was 50,000 and 40,000 respectively in WT and KO. In the NK cell-specific *Kcnj8* KO mice, there was a ∼50 % decrease of the most mature CD27^-^CD11b^+^ NK cells, with a corresponding increase in the immature CD27^+^CD11b^-^ cell population (**Figure 10C**).

**Figure 10:**
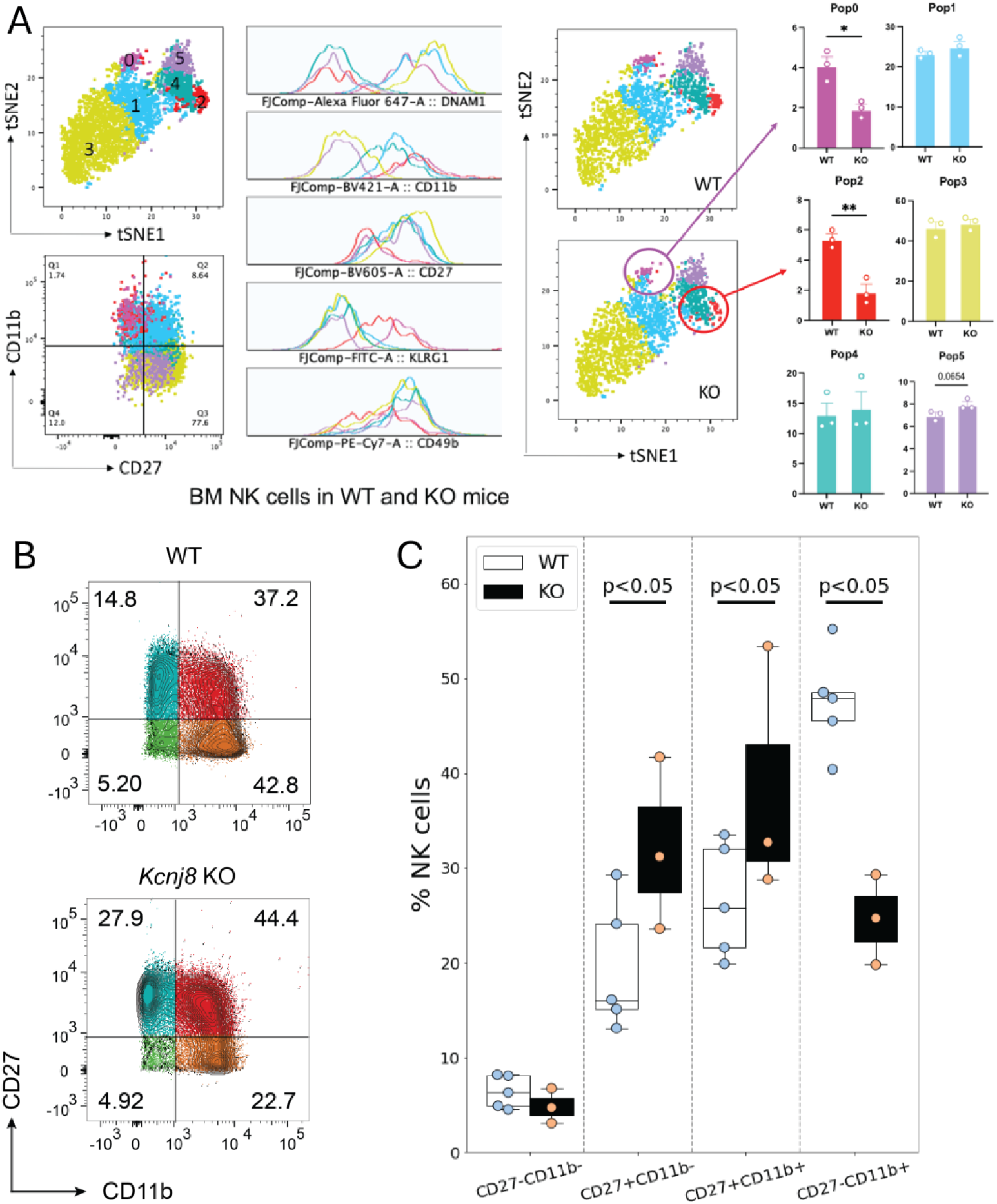
NK cell maturation in bone marrow and spleen of NK-specific *kcnj8* KO and WT mice. A) NK cells, isolated from bone marrow, were subjected to flow cytometry. Data of WT and constitutive NK-cell specific *Kcnj8* deficient mice are contrasted. A tSNE plot of differentially expressed proteins identified six populations of NK cells with differential protein expression (population 0 – population 5). NK maturation markers in each population is shown. WT is shown on the top and KO is the bottom. The frequency of each of the NK cell populations are compared between WT and KO mice. *p<0.05; *p<0.001 using a Student’s t-test. B). Isolated NK cells from WT mice, or mice with tamoxifen-induced NK-cell specific *Kcnj8*-deficiency, were subjected to flow cytometry. NK cells were selected based on expression of NK1.1 and NKp46. Cells were further selected based on expression of CD27 and CD11b. C) Summary data from 3-4 separate mice in each group is depicted as bar graphs. *p<0.05 with 2W-ANOVA, followed by a Dunnett’s t-test.

## Discussion

In this study, we found that *Kcnj8* is highly expressed in NK cell subsets of the mouse spleen and uterus. Of note, *Kcnj8* expression is highest in mature, educated NK cells. With patch clamping we recorded ionic currents in NK cells that are inhibited by the Kir6.1 blocker PNU-37883A. Using several approaches, we did not observe a direct effect of K_ATP_ channel blockers or *Kcnj8* deficiency on degranulation of NK cells – both *in vitro* and *in vivo*. We similarly did not observe an effect of *Kcnj8 in vivo* using a lymphoma cell tumor model. Pharmacological modulators of K_ATP_ channels had no effect on store-operated Ca^2+^ entry into NK cells. Transcriptomics shows that *Kcnj8* is highly expressed in mature CD27-/CD11b+ NK cells. Within this group, differentially expressed genes in the NK cell-specific KO mice include those with known roles in NK cell development. Indeed, we found that the developmental trajectory to be significantly impaired in both bone marrow and spleen of mice deficient of *Kcnj8* in NK cells. Overall, these data point to a previously unexplored cell-

### Kir6.1 as a K_ATP_ channel subunit in mouse NK cells

We found high expression of the *Kcnj8* K_ATP_ channel subunit in mouse NK cell subsets, which leads to the question whether K_ATP_ channels are involved in immunity. Indeed, there are indications in the literature that K_ATP_ channels may have a role in immunity. Patients with diabetes mellitus are more prone to bacterial sepsis. Glibenclamide, which is an anti-diabetic sulfonylurea (a K_ATP_ channel blocker), mitigates the secretion of proinflammatory cytokine production in polymorphonuclear neutrophils from diabetic patients, which might contribute to protection to bacterial infection (45). A glibenclamide-associated benefit was also observed with human melioidosis, which is correlated with an anti-inflammatory effect of the drug on the immune system (46). In mice, Kir6.1 (*Kcnj8*) deficiency has an exaggerated susceptibility to challenge with lipopolysaccharide (LPS) and survival is greatly impaired (47). A role for K_ATP_ channels in anti-viral pathways in *Drosophila* has also been noted (48, 49). A random mutagenesis screen in mice identified a mutation (named *mayday*) that caused a profound susceptibility to infection by mouse cytomegalovirus (MCMV) with decreased peak cytokine responses after inoculation. The *mayday* mice also had a ∼20,000-fold sensitization to LPS, and were hypersensitive to the lethal effects of poly(I.C) and CpG DNA (48). The *mayday* mutation was mapped to a deletion of *Kcnj8*, essentially leading to a knockout of *Kcnj8*. Interestingly, the *mayday* mice succumbed to infection by Listeria monocytogenes administered at doses sublethal to wild-type mice, but not to infection by vesicular stomatitis virus (VSV). The marked susceptibility in *mayday* mice to MCMV appears, however, not to be mediated by cells of hematopoietic origin, since mice still succumbed to infection after bone marrow transplantation. The protection of Kir6.1 against viral infection noted in this study may therefore reside in one or more tissues of extrahematopoietic origin. A caveat is that *Kcnj8* deficient mice succumb readily from cardiac arrhythmias during even mild stresses (50), and the high mortality of the *MayDay* mutants may be related to the stresses of irradiation and bone marrow transplantation.

We were able to record ion channels with pharmacological profile that matches that of K_ATP_ channels. The currents were insensitive to the K_ATP_ channel opener pinacidil. The lack of effect of pinacidil on the currents can be explained if the K_ATP_ channels were constitutively open in these cells under our experimental conditions. Moreover, the Kir6.1 blocker PNU-37883A inhibited ionic current magnitude, but only in cells where the current amplitude was large. The interpretation of the patch clamp experiments is complicated by the finding that only a subset of NK cells expresses *Kcnj8*. Selecting specific NK cell subsets (e.g. with CD11b labeling) presents additional challenges since the success of the patch clamp technique is significantly diminished by cell sorting and the presence of surface antibodies.

Functional K_ATP_ channels are formed by co-assembly of Kir6.1 (or Kir6.2) with one of the sulphonylurea receptors, SUR1 or SUR2 (22). Although mouse NK cells robustly express *Kcnj8*, the expression of *Abcc8* (SUR1) is absent and *Abcc9* (SUR2) mRNA expression is very low. Even though we can measure PNU-37883A sensitive currents in mouse NK cells with patch clamp approaches, we are open to the possibilities that PNU-37883A may not be completely specific against Kir6.1, and that Kir6.1 may have non-channel roles to regulate NK cell maturation and education. There is ample precedent for ion channel subunits that have functions beyond their role as ion channels. For example for Kv2 channels to participate in the formation of endoplasmic reticulum/plasma membrane junctions (51), the BKCa channel subunit not only forms Ca^2+^-activated K^+^ channels but also has roles in cell signaling and gene expression regulation (52), and Navβ subunits of voltage-gated Na^+^ channels that are involved in cell adhesion, particularly during development, and can influence cell migration and neurite outgrowth (53). Experiments to determine possible non-channel roles for Kir6.1 in NK cells are ongoing.

### Species differences in NK cell function and channel expression

NK cells from different species express different proteins and markers to perform the same functions (5). It would not be surprising, therefore, if ion channels are also differentially expressed (and have different roles) in various species. There is some controversy, for example, in the role of CRAC channels in degranulation and cytotoxicity. Mice deficient in CRAC channels (double knockout of STIM1 and STIM2) show robust NK cell degranulation/cytotoxicity and *in vivo* tumor rejection (54) – despite the established role for STIM/ORAI and SOCE in degranulation and cytotoxicity in human NK cells (21). Interventions designed to increase intracellular Ca^2+^ (ionomycin) increased INF-γ secretion, but not degranulation, whereas PKC activation with phorbol esters had the opposite effect (54). By contrast, unpublished data suggest that that NK cells from ORAI1-R93W knock-in mice (equivalent to the loss-of function R91W mutation in humans) are in fact deficient in target-cell induced degranulation and cytokine production [these data were mentioned in a previous publication (21)]. This issue remains to be resolved.

Differences in expression and function of other ion channels and transporters between mice and humans also remain to be elucidated. The negative membrane potential of most mammalian cells is determined by K^+^ channels. In NK cells, a role of the resting potential is illustrated by the finding that depolarizing NK cells with a high K^+^ results in partial inhibition of lysis (55). Moreover, a negative membrane potential is needed to maintain Ca^2+^ oscillations in NK cells following cross-linking with anti-CD16 (56). Early studies have revealed that K^+^ channels are indeed involved in the adhesion of NK cells to target cells and are crucial for lysis of target cells by NK cells from human peripheral blood (11, 12). In these studies, K^+^ currents were recorded from the NK cells. These channels were blocked by 4-aminopyridine, quinidine and by the traditional Ca^2+^ channel blockers verapamil and Cd^2+^. Moreover, the K^+^ channel blockers quinidine, Cd^2+^ and 4-AP, but not TEA, inhibited NK cell mediated killing of target cells (10, 11, 55).

At the molecular level, the expression and functional relevance of K^+^ channels is best characterized in human T lymphocytes, which express channels subunits such as KCa3.1 and Kv1.3 (9). There is evidence that these two channels may also be present in human NK cells. Indeed, expression of the voltage-gated Kv1.3 and the Ca^2+^-activated KCa3.1 K^+^ channel was confirmed in human peripheral blood NK cells (15). Interestingly, after activation of the NK cells by mitogens or tumor cells, adherent NK cells were found to preferentially upregulate KCa3.1, whereas non-adherent NK cells preferentially upregulate Kv1.3. Blockers of Kv1.3 mitigated the proliferation and degranulation of non-adherent NK cells, with minimal effects on the adherent NK cells. There is also evidence for a role of a K^+^ channel from a different class, namely the two-pore domain family. Specifically, K2p5.1 (also named TASK2) is expressed in human peripheral blood NK cells (17). An interesting observation in the latter study was K^+^ channel expression is distinctly regulated during maturation as NK cells progress from the immature CD56-/CD16- cells, to CD56^bright^/CD16-, CD56^bright^/CD16^dim^ and the most cytotoxic CD56^dim^/CD16^+^ subpopulations. K2p5.1 was downregulated during maturation through these cell stages, whereas Kv1.3 was upregulated. Thus, the complement of K^+^ channels, and their roles in NK cell function, may depend on the state of the human NK cell.

An avenue that needs to be explored is whether Kir6.1 is involved in NK cell Ca^2+^ signaling other than SOCE. For example, PI(3,5)P_2_-regulated Ca^2+^ release from acidic stores (via TRPML1) increases granzyme B expression and the functional potential of mouse NK cells, thereby mimicking the educated state (57), which in C57BL6 mice is marked by expression of Ly49C/I and NKG2A receptors (58). When Ca^2+^ is release across the membranes of intracellular stores, K^+^ fluxes through K^+^ channels are needed to maintain flux balance and electroneutrality (59). We previously found that Kir6.1/SUR2 containing K_ATP_ channels can localize to late endosomes and lysosomes (60) and the possibility is therefore raised that Kir6.1/SUR2 contributes Ca^2+^ release from acidic stores, thereby regulating NK cell education.

Our transcriptomics analysis of the mature CD27-/CD11b+ cells indicated that a) this NK cell population expresses the highest *Kcnj8* mRNA levels by far, and b) that downregulated genes with *Kcnj8* deficiency include *Il7r* (interleukin 7 receptor or IL7R), *Ltb* (Lymphotoxin B), and *kit* (KIT proto-oncogene receptor tyrosine kinase; c-Kit). All of these genes have critical roles in NK cell development (42–44). Indeed, we found that the NK cell maturity profile is significantly impacted in the bone marrow and spleen of mice with NK cell-specific *Kcnj8* deficiency. The transcriptomics dataset can be further explored in future experiment since several other genes are either up- or downregulated by NK cell-specific *Kcnj8* deficiency in the CD27-/CD11b+ cells (**Table 1**). Repression of the Notch1 target gene *Dtx1*, for example, prevents NK-cell differentiation (61). *Tigit* (T cell immunoreceptor with Ig and ITIM domains), which is upregulated with *Kcnj8* deficiency. The interaction between *Kcnj8* and *Tigit* might be an attractive candidate to explore in future experiments given the latter gene’s roles in in NK cell education, function, activation, functional heterogeneity, exhaustion, and antitumor responses (62–66).

## Grants

This work was supported by NIH R01HL146514 and R01HL148609 (W.A.C), the Wellcome Trust 200841/Z/16/Z (F.C.) and R01 AI097302, R01 AI130143, and R01 DE027981 to S.F., R03 TR004157 and R03 TR004459 to W.A.C. and S.F. A.S. is funded by the Medical Research Council grant MR/P001092/1. The NYUMC Experimental Pathology [RRID:SCR_017928] is part of the Division of Advanced Research Technologies and has received funds by the following grants: NIH/NCI 5 P30CA16087 and S10 OD021747.

## Acknowledgements

The computational requirements for this work were supported in part by the NYU Langone High Performance Computing (HPC) Core an d the Cytometry and Cell Sorting Laboratory (RRID: SCR_019179) resources and personnel. NIHR Cambridge BRC Cell Phenotyping Hub for their help with flow cytometry. Delia Hawkes is acknowledged for assistance with animal husbandry, mice breeding, and organ collections. We thank Cambridge Genomic services for uNK cells bulk RNA seq data analysis

## Conflict of Interest

S. F. is a scientific cofounder and consultant of CalciMedica. The other authors declare no conflicts of interest.

## References

1. Ikeda, H., Old, L. J., and Schreiber, R. D. (2002) The roles of IFN gamma in protection against tumor development and cancer immunoediting. Cytokine Growth Factor Rev. 13, 95–109

2. Wang, F., Qualls, A. E., Marques-Fernandez, L., and Colucci, F. (2021) Biology and pathology of the uterine microenvironment and its natural killer cells. Cell. Mol. Immunol. 18, 2101–2113

3. Abel, A. M., Yang, C., Thakar, M. S., and Malarkannan, S. (2018) Natural Killer Cells: Development, Maturation, and Clinical Utilization. Front Immunol 9, 1869

4. Colucci, F., and Traherne, J. (2017) Killer-cell immunoglobulin-like receptors on the cusp of modern immunogenetics. Immunology 152, 556–561

5. Colucci, F., Di Santo, J. P., and Leibson, P. J. (2002) Natural killer cell activation in mice and men: different triggers for similar weapons? Nat. Immunol. 3, 807–813

6. Stebbins, C. C., Watzl, C., Billadeau, D. D., Leibson, P. J., Burshtyn, D. N., and Long, E. O. (2003) Vav1 dephosphorylation by the tyrosine phosphatase SHP-1 as a mechanism for inhibition of cellular cytotoxicity. Mol. Cell. Biol. 23, 6291–6299

7. Anfossi, N., Andre, P., Guia, S., Falk, C. S., Roetynck, S., Stewart, C. A., Breso, V., Frassati, C., Reviron, D., Middleton, D., Romagne, F., Ugolini, S., and Vivier, E. (2006) Human NK cell education by inhibitory receptors for MHC class I. Immunity 25, 331–342

8. Feske, S., Concepcion, A. R., and Coetzee, W. A. (2019) Eye on ion channels in immune cells. Sci. Signal. 12

9. Feske, S., Wulff, H., and Skolnik, E. Y. (2015) Ion channels in innate and adaptive immunity. Annu. Rev. Immunol. 33, 291–353

10. Huwyler, T., Hirt, A., Felix, D., and Morell, A. (1985) Effect of cations and cation channel blockers on human natural killer cells. Int. J. Immunopharmacol. 7, 573–576

11. Schlichter, L., Sidell, N., and Hagiwara, S. (1986) Potassium channels mediate killing by human natural killer cells. Proc. Natl. Acad. Sci. U. S. A. 83, 451–455

12. Sidell, N., Schlichter, L. C., Wright, S. C., Hagiwara, S., and Golub, S. H. (1986) Potassium channels in human NK cells are involved in discrete stages of the killing process. J. Immunol. 137, 1650–1658

13. Decoursey, T. E., Chandy, K. G., Gupta, S., and Cahalan, M. D. (1987) Two types of potassium channels in murine T lymphocytes. J. Gen. Physiol. 89, 379–404

14. Lewis, R. S., and Cahalan, M. D. (1995) Potassium and calcium channels in lymphocytes. Annu. Rev. Immunol. 13, 623–653

15. Koshy, S., Wu, D., Hu, X., Tajhya, R. B., Huq, R., Khan, F. S., Pennington, M. W., Wulff, H., Yotnda, P., and Beeton, C. (2013) Blocking KCa3.1 channels increases tumor cell killing by a subpopulation of human natural killer lymphocytes. PLoS ONE 8, e76740

16. Olivas-Aguirre, M., Cruz-Aguilar, L. H., Pottosin, I., and Dobrovinskaya, O. (2023) Reduction of Ca(2+) Entry by a Specific Block of KCa3.1 Channels Optimizes Cytotoxic Activity of NK Cells against T-ALL Jurkat Cells. Cells 12

17. Schulte-Mecklenbeck, A., Bittner, S., Ehling, P., Doring, F., Wischmeyer, E., Breuer, J., Herrmann, A. M., Wiendl, H., Meuth, S. G., and Gross, C. C. (2015) The two-pore domain K2 P channel TASK2 drives human NK-cell proliferation and cytolytic function. Eur. J. Immunol. 45, 2602–2614

18. Grandclement, C., Pick, H., Vogel, H., and Held, W. (2016) NK Cells Respond to Haptens by the Activation of Calcium Permeable Plasma Membrane Channels. PLoS ONE 11, e0151031

19. Nguyen, T., Johnston, S., Clarke, L., Smith, P., Staines, D., and Marshall-Gradisnik, S. (2017) Impaired calcium mobilization in natural killer cells from chronic fatigue syndrome/myalgic encephalomyelitis patients is associated with transient receptor potential melastatin 3 ion channels. Clin. Exp. Immunol. 187, 284–293

20. Balinas, C., Cabanas, H., Staines, D., and Marshall-Gradisnik, S. (2019) Identification and characterisation of transient receptor potential melastatin 2 and CD38 channels on natural killer cells using the novel application of flow cytometry. BMC Immunol. 20, 14

21. Maul-Pavicic, A., Chiang, S. C., Rensing-Ehl, A., Jessen, B., Fauriat, C., Wood, S. M., Sjoqvist, S., Hufnagel, M., Schulze, I., Bass, T., Schamel, W. W., Fuchs, S., Pircher, H., McCarl, C. A., Mikoshiba, K., Schwarz, K., Feske, S., Bryceson, Y. T., and Ehl, S. (2011) ORAI1-mediated calcium influx is required for human cytotoxic lymphocyte degranulation and target cell lysis. Proc. Natl. Acad. Sci. U. S. A. 108, 3324–3329

22. Foster, M. N., and Coetzee, W. A. (2016) KATP Channels in the Cardiovascular System. Physiol. Rev. 96, 177–252

23. Tinker, A., Aziz, Q., Li, Y., and Specterman, M. (2018) ATP-Sensitive Potassium Channels and Their Physiological and Pathophysiological Roles. Comprehensive Physiology 8, 1463–1511

24. Ashcroft, F. M., Kakei, M., Kelly, R. P., and Sutton, R. (1987) ATP-sensitive K^+^ channels in human isolated pancreatic B-cells. FEBS Lett. 215, 9–12

25. Martin, G. M., Yoshioka, C., Rex, E. A., Fay, J. F., Xie, Q., Whorton, M. R., Chen, J. Z., and Shyng, S. L. (2017) Cryo-EM structure of the ATP-sensitive potassium channel illuminates mechanisms of assembly and gating. eLife 6

26. Sung, M. W., Yang, Z., Driggers, C. M., Patton, B. L., Mostofian, B., Russo, J. D., Zuckerman, D. M., and Shyng, S. L. (2021) Vascular KATP channel structural dynamics reveal regulatory mechanism by Mg-nucleotides. Proc. Natl. Acad. Sci. U. S. A. 118

27. Nabekura, T., and Lanier, L. L. (2016) Tracking the fate of antigen-specific versus cytokine-activated natural killer cells after cytomegalovirus infection. J. Exp. Med. 213, 2745–2758

28. Narni-Mancinelli, E., Chaix, J., Fenis, A., Kerdiles, Y. M., Yessaad, N., Reynders, A., Gregoire, C., Luche, H., Ugolini, S., Tomasello, E., Walzer, T., and Vivier, E. (2011) Fate mapping analysis of lymphoid cells expressing the NKp46 cell surface receptor. Proc. Natl. Acad. Sci. U. S. A. 108, 18324–18329

29. Hao, Y., Hao, S., Andersen-Nissen, E., Mauck, W. M., 3rd, Zheng, S., Butler, A., Lee, M. J., Wilk, A. J., Darby, C., Zager, M., Hoffman, P., Stoeckius, M., Papalexi, E., Mimitou, E. P., Jain, J., Srivastava, A., Stuart, T., Fleming, L. M., Yeung, B., Rogers, A. J., McElrath, J. M., Blish, C. A., Gottardo, R., Smibert, P., and Satija, R. (2021) Integrated analysis of multimodal single-cell data. Cell 184, 3573–3587 e3529

30. Xie, Z., Bailey, A., Kuleshov, M. V., Clarke, D. J. B., Evangelista, J. E., Jenkins, S. L., Lachmann, A., Wojciechowicz, M. L., Kropiwnicki, E., Jagodnik, K. M., Jeon, M., and Ma’ayan, A. (2021) Gene Set Knowledge Discovery with Enrichr. Curr Protoc 1, e90

31. Depierreux, D. M., Seshadri, E., Shmeleva, E. V., Kieckbusch, J., Hawkes, D. A., and Colucci, F. (2021) Isolation of Uterine Innate Lymphoid Cells for Analysis by Flow Cytometry. J. Vis. Exp.

32. Depierreux, D. M., Smith, G. L., and Ferguson, B. J. (2023) Transcriptional reprogramming of natural killer cells by vaccinia virus shows both distinct and conserved features with mCMV. Front Immunol 14, 1093381

33. Zhang, J., Le Gras, S., Pouxvielh, K., Faure, F., Fallone, L., Kern, N., Moreews, M., Mathieu, A. L., Schneider, R., Marliac, Q., Jung, M., Berton, A., Hayek, S., Vidalain, P. O., Marcais, A., Dodard, G., Dejean, A., Brossay, L., Ghavi-Helm, Y., and Walzer, T. (2021) Sequential actions of EOMES and T-BET promote stepwise maturation of natural killer cells. Nat. Commun. 12, 5446

34. Wu, Z., Park, S., Lau, C. M., Zhong, Y., Sheppard, S., Sun, J. C., Das, J., Altan-Bonnet, G., and Hsu, K. C. (2021) Dynamic variability in SHP-1 abundance determines natural killer cell responsiveness. Sci. Signal. 14, eabe5380

35. Schmied, L., Luu, T. T., Sondergaard, J. N., Hald, S. H., Meinke, S., Mohammad, D. K., Singh, S. B., Mayer, C., Perinetti Casoni, G., Chrobok, M., Schlums, H., Rota, G., Truong, H. M., Westerberg, L. S., Guarda, G., Alici, E., Wagner, A. K., Kadri, N., Bryceson, Y. T., Saeed, M. B., and Hoglund, P. (2023) SHP-1 localization to the activating immune synapse promotes NK cell tolerance in MHC class I deficiency. Sci. Signal. 16, eabq0752

36. Kovalev, H., Quayle, J. M., Kamishima, T., and Lodwick, D. (2004) Molecular analysis of the subtype-selective inhibition of cloned KATP channels by PNU-37883A. Br. J. Pharmacol. 141, 867–873

37. Cui, Y., Tinker, A., and Clapp, L. H. (2003) Different molecular sites of action for the KATP channel inhibitors, PNU-99963 and PNU-37883A. Br. J. Pharmacol. 139, 122–128

38. Shabrish, S., Gupta, M., and Madkaikar, M. (2016) A Modified NK Cell Degranulation Assay Applicable for Routine Evaluation of NK Cell Function. J Immunol Res 2016, 3769590

39. Diefenbach, A., Jensen, E. R., Jamieson, A. M., and Raulet, D. H. (2001) Rae1 and H60 ligands of the NKG2D receptor stimulate tumour immunity. Nature 413, 165–171

40. Hayakawa, Y., Kelly, J. M., Westwood, J. A., Darcy, P. K., Diefenbach, A., Raulet, D., and Smyth, M. J. (2002) Cutting edge: tumor rejection mediated by NKG2D receptor-ligand interaction is dependent upon perforin. J. Immunol. 169, 5377–5381

41. Prakriya, M., and Lewis, R. S. (2015) Store-Operated Calcium Channels. Physiol. Rev. 95, 1383–1436

42. Williams, N. S., Klem, J., Puzanov, I. J., Sivakumar, P. V., Bennett, M., and Kumar, V. (1999) Differentiation of NK1.1+, Ly49+ NK cells from flt3+ multipotent marrow progenitor cells. J. Immunol. 163, 2648–2656

43. Iizuka, K., Chaplin, D. D., Wang, Y., Wu, Q., Pegg, L. E., Yokoyama, W. M., and Fu, Y. X. (1999) Requirement for membrane lymphotoxin in natural killer cell development. Proc. Natl. Acad. Sci. U. S. A. 96, 6336–6340

44. Colucci, F., and Di Santo, J. P. (2000) The receptor tyrosine kinase c-kit provides a critical signal for survival, expansion, and maturation of mouse natural killer cells. Blood 95, 984–991

45. Kewcharoenwong, C., Rinchai, D., Utispan, K., Suwannasaen, D., Bancroft, G. J., Ato, M., and Lertmemongkolchai, G. (2013) Glibenclamide reduces pro-inflammatory cytokine production by neutrophils of diabetes patients in response to bacterial infection. Scientific reports 3, 3363

46. Koh, G. C., Maude, R. R., Schreiber, M. F., Limmathurotsakul, D., Wiersinga, W. J., Wuthiekanun, V., Lee, S. J., Mahavanakul, W., Chaowagul, W., Chierakul, W., White, N. J., van der Poll, T., Day, N. P., Dougan, G., and Peacock, S. J. (2011) Glyburide is anti-inflammatory and associated with reduced mortality in melioidosis. Clin. Infect. Dis. 52, 717–725

47. Kane, G. C., Lam, C. F., O’Cochlain, F., Hodgson, D. M., Reyes, S., Liu, X. K., Miki, T., Seino, S., Katusic, Z. S., and Terzic, A. (2006) Gene knockout of the KCNJ8-encoded Kir6.1 K(ATP) channel imparts fatal susceptibility to endotoxemia. FASEB J. 20, 2271–2280

48. Croker, B., Crozat, K., Berger, M., Xia, Y., Sovath, S., Schaffer, L., Eleftherianos, I., Imler, J. L., and Beutler, B. (2007) ATP-sensitive potassium channels mediate survival during infection in mammals and insects. Nat. Genet. 39, 1453–1460

49. Eleftherianos, I., Won, S., Chtarbanova, S., Squiban, B., Ocorr, K., Bodmer, R., Beutler, B., Hoffmann, J. A., and Imler, J. L. (2011) ATP-sensitive potassium channel (K(ATP))-dependent regulation of cardiotropic viral infections. Proc. Natl. Acad. Sci. U. S. A. 108, 12024–12029

50. Miki, T., Suzuki, M., Shibasaki, T., Uemura, H., Sato, T., Yamaguchi, K., Koseki, H., Iwanaga, T., Nakaya, H., and Seino, S. (2002) Mouse model of Prinzmetal angina by disruption of the inward rectifier Kir6.1. Nat. Med. 8, 466–472

51. Johnson, B., Leek, A. N., Sole, L., Maverick, E. E., Levine, T. P., and Tamkun, M. M. (2018) Kv2 potassium channels form endoplasmic reticulum/plasma membrane junctions via interaction with VAPA and VAPB. Proc. Natl. Acad. Sci. U. S. A.

52. Toro, L., Li, M., Zhang, Z., Singh, H., Wu, Y., and Stefani, E. (2014) MaxiK channel and cell signalling. Pflugers Arch. 466, 875–886

53. O’Malley, H. A., and Isom, L. L. (2015) Sodium channel beta subunits: emerging targets in channelopathies. Annu. Rev. Physiol. 77, 481–504

54. Freund-Brown, J., Choa, R., Singh, B. K., Robertson, T. F., Ferry, G. M., Viver, E., Bassiri, H., Burkhardt, J. K., and Kambayashi, T. (2017) Cutting Edge: Murine NK Cells Degranulate and Retain Cytotoxic Function without Store-Operated Calcium Entry. J. Immunol.

55. Ng, J., Fredholm, B. B., and Jondal, M. (1987) Studies on the calcium dependence of human NK cell killing. Biochem. Pharmacol. 36, 3943–3949

56. Hess, S. D., Oortgiesen, M., and Cahalan, M. D. (1993) Calcium oscillations in human T and natural killer cells depend upon membrane potential and calcium influx. J. Immunol. 150, 2620–2633

57. Goodridge, J. P., Jacobs, B., Saetersmoen, M. L., Clement, D., Hammer, Q., Clancy, T., Skarpen, E., Brech, A., Landskron, J., Grimm, C., Pfefferle, A., Meza-Zepeda, L., Lorenz, S., Wiiger, M. T., Louch, W. E., Ask, E. H., Liu, L. L., Oei, V. Y. S., Kjallquist, U., Linnarsson, S., Patel, S., Tasken, K., Stenmark, H., and Malmberg, K. J. (2019) Remodeling of secretory lysosomes during education tunes functional potential in NK cells. Nat. Commun. 10, 514

58. Clement, D., Goodridge, J. P., Grimm, C., Patel, S., and Malmberg, K. J. (2020) TRP Channels as Interior Designers: Remodeling the Endolysosomal Compartment in Natural Killer Cells. Front Immunol 11, 753

59. Kuum, M., Veksler, V., and Kaasik, A. (2015) Potassium fluxes across the endoplasmic reticulum and their role in endoplasmic reticulum calcium homeostasis. Cell Calcium 58, 79–85

60. Bao, L., Hadjiolova, K., Coetzee, W. A., and Rindler, M. J. (2011) Endosomal KATP channels as a reservoir after myocardial ischemia: a role for SUR2 subunits. Am. J. Physiol. Heart Circ. Physiol. 300, H262–270

61. Van de Walle, I., Dolens, A. C., Durinck, K., De Mulder, K., Van Loocke, W., Damle, S., Waegemans, E., De Medts, J., Velghe, I., De Smedt, M., Vandekerckhove, B., Kerre, T., Plum, J., Leclercq, G., Rothenberg, E. V., Van Vlierberghe, P., Speleman, F., and Taghon, T. (2016) GATA3 induces human T-cell commitment by restraining Notch activity and repressing NK-cell fate. Nat. Commun. 7, 11171

62. Ziegler, A. E., Fittje, P., Muller, L. M., Ahrenstorf, A. E., Hagemann, K., Hagen, S. H., Hess, L. U., Niehrs, A., Poch, T., Ravichandran, G., Lobl, S. M., Padoan, B., Brias, S., Hennesen, J., Richard, M., Richert, L., Peine, S., Oldhafer, K. J., Fischer, L., Schramm, C., Martrus, G., Bunders, M. J., Altfeld, M., and Lunemann, S. (2023) The co-inhibitory receptor TIGIT regulates NK cell function and is upregulated in human intrahepatic CD56(bright) NK cells. Front Immunol 14, 1117320

63. Wang, F., Hou, H., Wu, S., Tang, Q., Liu, W., Huang, M., Yin, B., Huang, J., Mao, L., Lu, Y., and Sun, Z. (2015) TIGIT expression levels on human NK cells correlate with functional heterogeneity among healthy individuals. Eur. J. Immunol. 45, 2886–2897

64. Zhang, Q., Bi, J., Zheng, X., Chen, Y., Wang, H., Wu, W., Wang, Z., Wu, Q., Peng, H., Wei, H., Sun, R., and Tian, Z. (2018) Blockade of the checkpoint receptor TIGIT prevents NK cell exhaustion and elicits potent anti-tumor immunity. Nat. Immunol. 19, 723–732

65. Hasan, M. F., Campbell, A. R., Croom-Perez, T. J., Oyer, J. L., Dieffenthaller, T. A., Robles-Carrillo, L. D., Cash, C. A., Eloriaga, J. E., Kumar, S., Andersen, B. W., Naeimi Kararoudi, M., Tullius, B. P., Lee, D. A., and Copik, A. J. (2023) Knockout of the inhibitory receptor TIGIT enhances the antitumor response of ex vivo expanded NK cells and prevents fratricide with therapeutic Fc-active TIGIT antibodies. J Immunother Cancer 11

66. He, Y., Peng, H., Sun, R., Wei, H., Ljunggren, H. G., Yokoyama, W. M., and Tian, Z. (2017) Contribution of inhibitory receptor TIGIT to NK cell education. J. Autoimmun. 81, 1–12

