## Supplemental Information for "Kir6.1, a component of an ATP-sensitive potassium channel, regulates natural killer cell development"

### Supplemental Figures

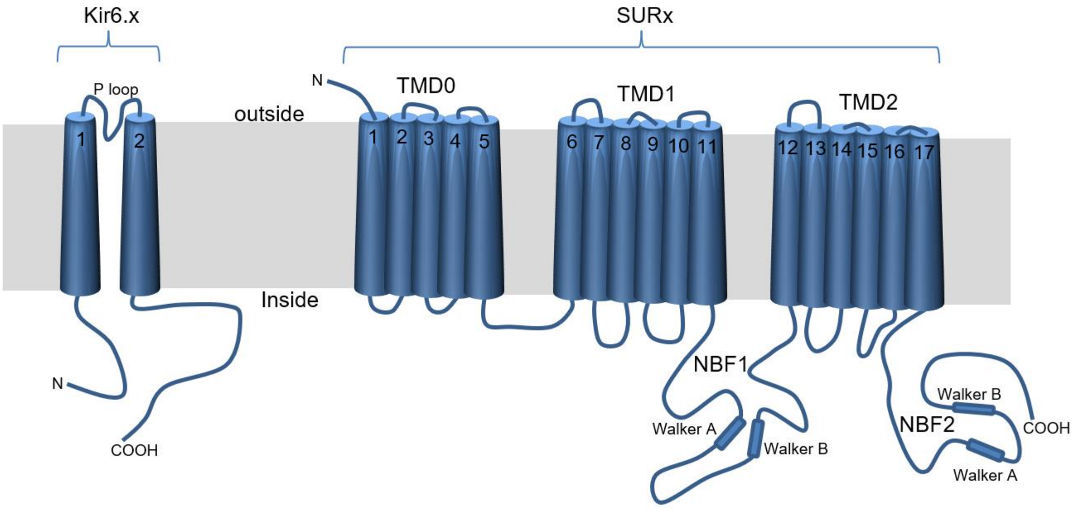


Show/hide columns

| No. | Exon / Intron | Start | End | Start Phase | End Phase | Length | Sequence |
|---|---|---|---|---|---|---|---|
| | 5' upstream sequence | | | | | | ..........cgagccttttaaagatgtgtgtgccggtctggcagcagaatccgggctac |
| 1 | ENSMUSE00001365548 | 142,517,340 | 142,516,910 | - | - | 431 | TGGGAACCGAGCTGGCTCGGCTGGTGCCTCCTCCCCCGGTGGTCCCTCCCTTCTGGCCGG<br>CTCCCTGCCTCCCCGGCTCCCCCGACTGGCCTCCGCCTCCACTTTCCCCGGCTCGCTCGG<br>AGGTCTGGCCCCAGCTAGGCGGCTGAGGAGCTGGCGGAGCATGGCCACTAGCACCTCTAT<br>CCCCAGCTCCTACCCCGCCCTCCCCGTGGCCCGGCAGCAGGAGGGGGGAAGCCAAACCCA<br>ACCCTCATCTTCCAGCCCTCCTCCCCGCGTGCACTATGGATCGCACCTTCTAGAGTCTTA<br>ACTCAGTTCTGGAGGACCAACATCCCCCGGATCTGCACTTCGGGAGGTCTCTGCTCCCGG<br>GATGCGAGAGCCTGGGACCAGCCCGCCCTGTGCGGAGCTCAAGCCGCAGCCGGGAGCGCA<br>AACCCGAGTCT |
| | Intron 1-2 | 142,516,909 | 142,516,180 | | | 730 | gtaagttgcatcgcgcctcccgggg..........cagtttttaataagcgtttctctag |
| 2 | ENSMUSE00000196744 | 142,516,179 | 142,515,732 | - | 2 | 448 | TCTAGGAGGACGCGTGTGGAGGAAAGGAGCCACAGGTTCAGGCAGGTGCATAGGCGGGCT<br>ATGGTGAAAGGAAGATGTTGGCCAGGAAGAGCATCATCCCGGAGGAGTATGTGCTGGCGC<br>GCATCGCAGCGGAGAACCTGCGCAAACCGCGCATCCGCGACCGTCTCCCCAAAGCCCGCT<br>TCATCGCCAAGAGCGGAGCCTGCAACCTGGCACACAAGAACATCCGAGAGCAAGGTCGCT<br>TCCTGCAGGACATCTTCACCACCTTGGTAGACCTGAAGTGGCGTCACACGCTGGTCATCT<br>TCACCATGTCCTTCCTCTGCAGCTGGCTGCTCTTCGCTATCATGTGGTGGCTGGTGGCCT<br>TCGCCCACGGGGACATCTATGCTTACATGGAGAAAGGCACCATGGAGAAGAGTGGCCTGG<br>AGTCCGCTGTCTGTGTGACCAATGTCAG |
| | Intron 2-3 | 142,515,731 | 142,512,232 | | | 3,500 | gtaagaatgaaatggagttaccagc..........tcatttctgatattggtccttacag |
| 3 | ENSMUSE00000369261 | 142,512,231 | 142,510,563 | 2 | - | 1,669 | GTCATTCACGTCTGCGTTTCTCTTCTCCATTGAGGTTCAAGTGACCATTGGGTTTGGAGG<br>GAGAATGATGACTGAGGAATGCCCTCTGGCCATCACGGTTTTGATTCTGCAGAACATCGT<br>GGGTCTGATCATCAACGCAGTCATGTTGGGCTGCATCTTCATGAAGACGGCGCAGGCCCA<br>CAGAAGGGCAGAGACGCTGATTTTCAGCCGCCATGCTGTGATTGCCGTCCGCAATGGCAA<br>GCTGTGCTTCATGTTCCGGGTGGGTGACCTGAGGAAGAGCATGATCATTAGCGCCTCGGT<br>GCGCATCCAGGTGGTCAAGAAAACCACGACGCCAGAAGGGGAGGTGGTGCCTATTCATCA<br>GCAGGACATTCCTGTTGATAATCCCATCGAGAGCAATAATATCTTCCTAGTGGCCCCATT<br>GATCATCTGCCACGTGATTGACAAGCGTAGCCCCCTGTATGATATCTCAGCAACTGACCT<br>TGCCAATCAAGACCTGGAGGTCATAGTGATTCTCGAGGGCGTGGTAGAAACCACAGGCAT<br>CACCACACAAGCACGGACCTCCTACATTGCCGAGGAGATCCAGTGGGGACACCGCTTCGT<br>GTCAATTGTGACTGAGGAGGAGGGCGTGTACTCTGTGGACTATTCCAAATTTGGTAACAC<br>GGTGAGAGTGGCTGCGCCAAGATGCAGTGCCCGGGAGCTGGATGAGAAGCCTTCCATCCT<br>GATTCAGACCCTCCAAAAGAGCGAACTGTCGCACCAGAATTCTCTGCGGAAGCGCAACTC<br>CATGAGGAGAAACAACTCCATGAGGAGAAACAACTCCATCAGGAGGAATAACTCTTCCCT<br>CATGGTGCCCAAGGTGCAGTTCATGACTCCAGAAGGAAACCAGTGTCCATCAGAATCATG<br>AGGGCAGGATGACCGGAGACAGTTACTTGTTGAGTCCTGATGACTGATAGCCCTGAACAG<br>TCACTGTGTCCTGATGACTGAGAGACAATCCGGAGACAGTTCATTGAGTTCCGATGATCA<br>AAATATTGCACTCATCACCAGTTCAGGGCTGGAGCACAGTATTCCTATCCTAATGCACTG<br>AGAAATATTAATATTTGAGACATTAAACTTCCTGTATTAATAAACAATAACACACAAACC<br>TGAGCTCTTTATTCTCCTCTCATCTTAAAATTCTGTTTTCTTCCCAGCAGTCTCTGTAGT<br>CAGGTCAGTTGGCCTGTGTTTTTGAGCTTCTGTCACTTAGCAGAGATGAGATCTTCTACC<br>CAAGCCCAGTTTCGTCATTCTTAGGTCCTGCACCCTCAAGTAAGGGCAGAGTAGAGAGAC<br>GTCCTGGGTTAACACGGTGTGTGATGTGGGGTTAACCCAGTGTGTGATATGGGGGTAACA<br>CGTTGTGTGATGCGGTAAAACATGGAAGCTGGGCAGCTGGGTAGCCTTTTATTCAGAGTC |

**Figure S1:** $K_{ATP}$ channel structure and genomic organization of *Kcnj8*. Top: The $K_{ATP}$ channel is composed of two types of subunits: four Kir6.x in combination with four SURx subunits. Kir6.1 is a 2 transmembrane (TM) protein with intracellular N- and C-termini. The pore that conducts $K^+$ across the membrane is formed by the TM domains and the $K^+$ selectivity filter is composed of sequences within the P-loop. Bottom: The *Kcnj8* gene (Kir6.1 or potassium inwardly rectifying channel, subfamily J, member 8) is located on mouse chromosome 6: 142,564,939-142,571,356 on the reverse strand (ENSMUSG00000030247). It contains three exons. Exon 2 was targeted for deletion by introducing loxP sites in the flanking intronic sequences. Deletion of exon 2 results in the loss of the Kir6.1 N-terminus the M1 TM domain and part of the P-loop, such that no functional channel can be formed when exon 2 is deleted.

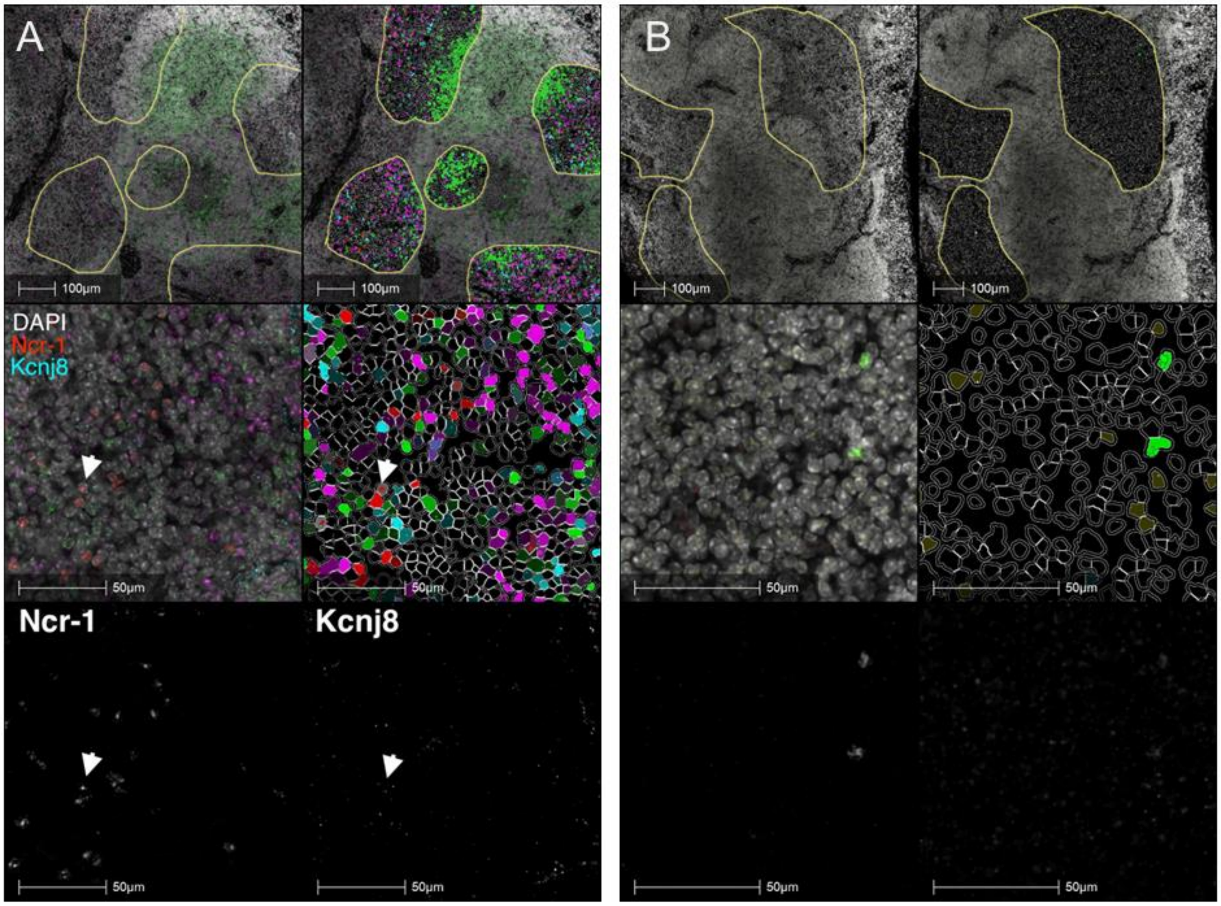


**Figure S2**: RNA Scope data of a mouse spleen. A) the top row represents images at low magnification, whereas the bottom two rows are images at higher resolution. Areas marked for cell detection analysis are demarcated with yellow lines. The top two rows of the left column show RNA Scope probe dots for *Ncr-1* (red) and *Kcnj8* (light blue), whereas DAPI staining is shown in gray. Cell segmentation was performed based on DAPI staining and cells are pseudo colored by predominant expression of *Ncr1* (red), *Kcnj8* (light blue), *CD3e* (green) and *Abgre1* (magenta). The bottom row depicts the RNA Scope dots (in the absence of other information) respectively for *Ncr-1* and *Kcnj8*. The arrows depict a cell that co-express dots for *Ncr-1* and *Kcnj8*, which is shown at higher resolution in **Figure 1**. B) A negative control experiment performed with probes against bacterial proteins.

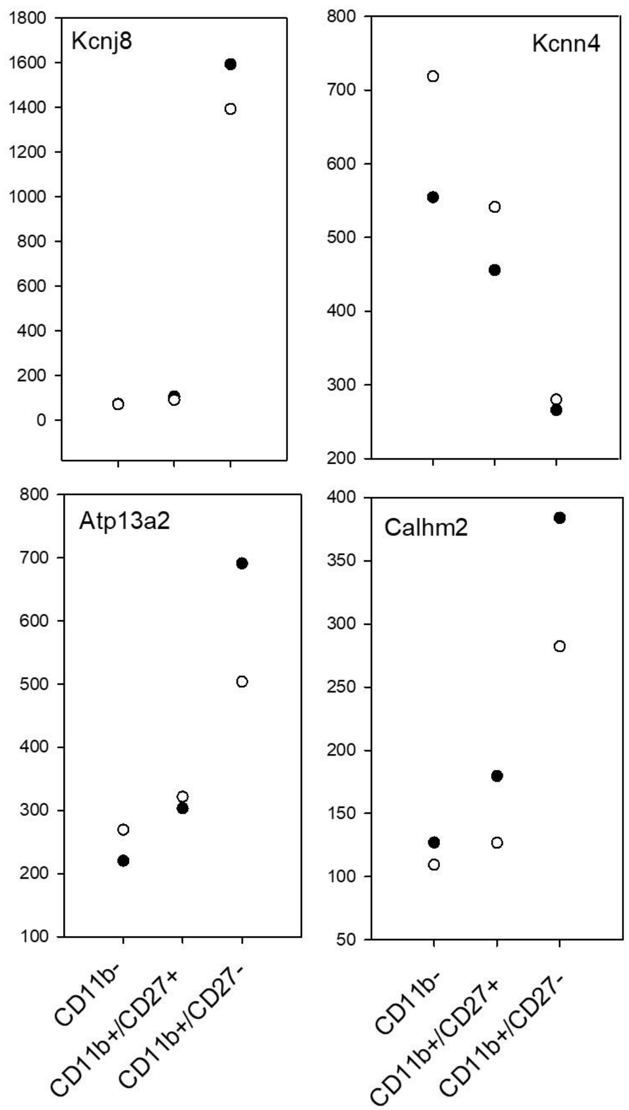


**Figure S3**: We examined RNA-seq profiling data obtained from NK1.1$^{+}$ cells isolated from the spleens of RAG$^{-/-}$ mice that were sorted into CD27$^{-}$CD11b$^{-}$, CD27$^{+}$CD11b$^{+}$ and CD27$^{-}$CD11b$^{+}$ subsets by flow cytometry (GEO GSE13229). Differential gene expression analysis with DEseq of the bulk RNA-seq data interestingly identified *Kcnj8* as the gene with the largest cause effect amongst all genes in the three groups ($p<0.00001$; $F=465.9$), with *Kcnj8* expressed specifically and at high levels in the most cytotoxic CD27$^{-}$CD11b$^{+}$ NK cells. Also shown are the only three other channel or transporter genes that are differentially expressed between the three groups in this dataset.

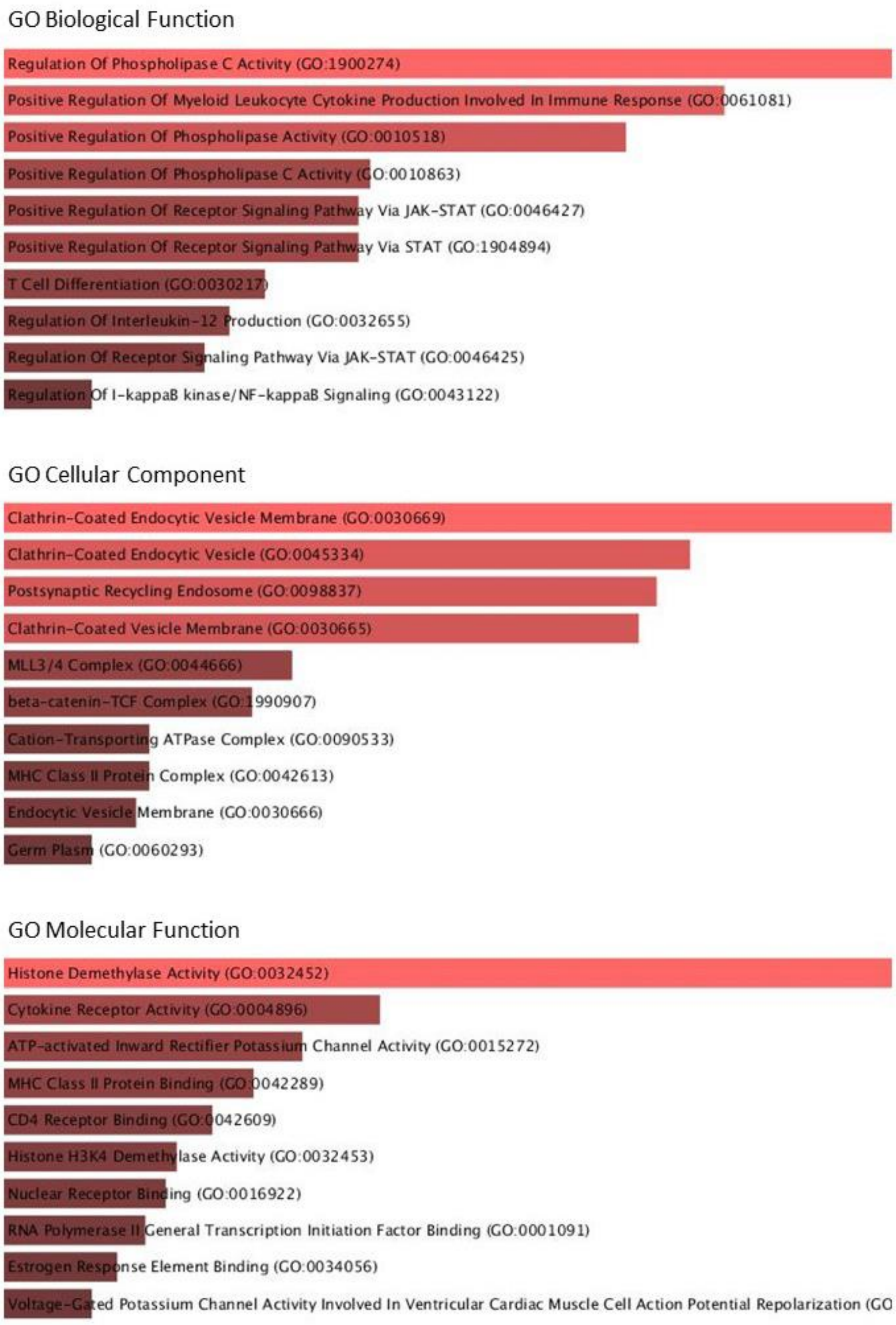


**Figure S4**: Gene ontology (GO) analysis performed with Enrichr. Bulk RNAseq was performed of mouse splenic CD27-/CD11b+ NK cells, and differential gene expression was calculated by comparing the WT and *Kcnj8* deficient genotypes. The GO analysis was performed with 29 genes that were differentially expressed.

NK cells Given that Kcnj8 expression is so much elevated in the CD27-/CD11b+ population, we performed a differential gene expression analysis in this population by comparing the WT and KO genotypes.

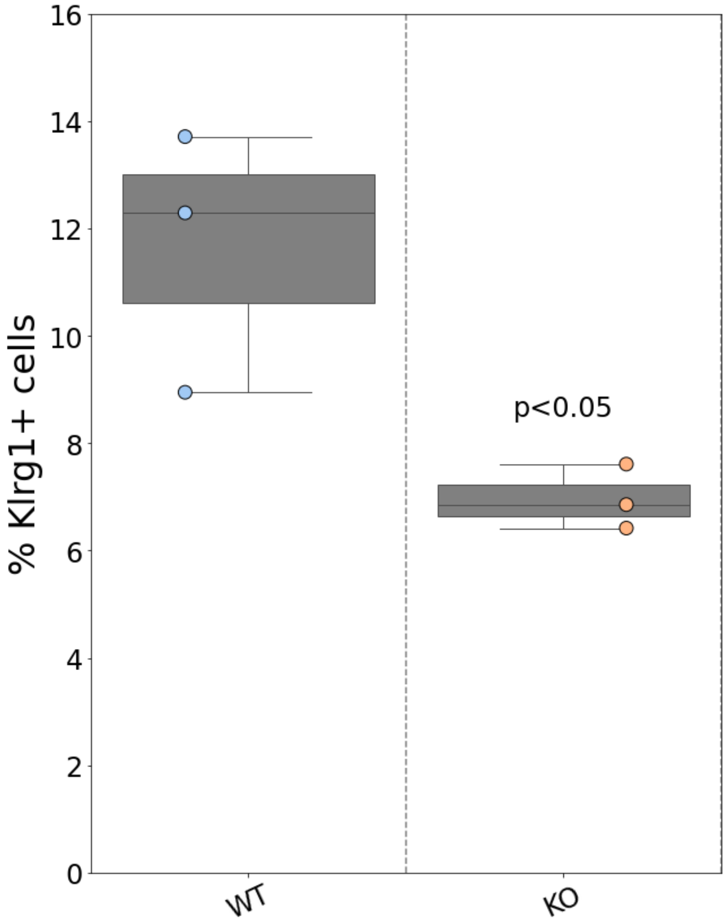


**Figure S5**. *Klrgt1* expression in bone marrow NK cells isolated from WT mice and NK cell-specific *kcnj8* KO mice. *p = 0.0321 with Student's t-test.

### SUPPLEMENTAL TABLES

**Table S1**: Antibodies used in flow cytometry and FACS

| Marker | Fluorochrome | Provider | Cat number | Clone | Dilution |
|---|---|---|---|---|---|
| CD3 | PE | BioLegend | 100206 | 17A2 | 1:100 |
| CD19 | PE | BD Bioscience | 152408 | 1D3 | 1:200 |
| CD4 | PE | BioLegend | 100408 | GK1.5 | 1:200 |
| CD8 | PE | BioLegend | 100708 | 53-6.7 | 1:300 |
| NK1.1 | PE | BioLegend | 108707 | PK136 | 1:200 |
| Ly49 C/I | R718 | BD Bioscience | 752258 | 5E6 | 1:50 |
| CD45 | FITC | BioLegend | 103108 | 30-F11 | 1:200 |
| CD11b | AF488 | BioLegend | 101219 | M1/70 | 1:200 |
| CD11b | APC | BioLegend | 101211 | M1/70 | 1:100 |
| NKp46 | APC | BioLegend | 137608 | 29A1.4 | 1:20 |
| NK1.1 | APC-Cy7 | BioLegend | 108723 | PK136 | 1:100 |
| CD27 | APC-Cy7 | BioLegend | 124225 | LG.3A10 | 1:100 |
| CD27 | AF647 | BioLegend | 124219 | LG.3A10 | 1:200 |
| CD107a | APC/Fire750 | BioLegend | 121633 | 1D4B | 1:100 |
| CD11b | BV785 | BioLegend | 101243 | M1/70 | 1:100 |
| NKG2AC/E | BV421 | BD | 740065 | 20d5 | 1:100 |
| CD335 (NKp46) | BV421 | BioLegend | 137612 | 29A1.4 | 1:50 |
| Zombie aqua | | BioLegend | 423101 | | 1:400 |
| Fc block | | BioLegend | 101320 | | 1:100 |

**Table S2**: Parameters used to detect the RNAscope dots

| Channel | Parameter | Value |
|---|---|---|
| | Class List | |
| | Classifier | |
| | Classifier Output Type | Mask |
| | Classify Registered | |
| | Image Zoom | 2 |
| | Membrane Masker | |
| | Membrane Masker Type | None |
| | Nuclear Masker | Nuclear_Segmentation_Mouse_spleen |
| | Nuclear Masker Type | AI Custom |
| | Phenotyper | |
| | Phenotyper Filter | |
| | Phenotyper Segmentation Type | |
| | nuclear_dye | DAPI |
| | num_probes | 4 |
| | detect_cells | TRUE |
| | use_nn | FALSE |
| | nuclear_contrast | 0.5 |
| | nuclear_min_intensity | 0.095 |

| | | | |
|---|---|---|---|
| | nuclear_segg_agg | 0.65 | |
| | nuclear_fill_holes | FALSE | |
| | nuclear_min_area | 11.3 | |
| | nuclear_max_area | 571.7 | |
| | nuclear_roundness | 0 | |
| | cyto_radius | 1 | |
| | localize_results | FALSE | |
| | min_plus1 | 1 | |
| | min_plus2 | 2 | |
| | min_plus3 | 3 | |
| | min_plus4 | 4 | |
| | num_phenotypes | 7 | |
| | verbose | TRUE | |
| | FISH Probe 1 | Ncr1-C2 (Opal 520) | |
| | FISH Probe 2 | Kcnj8-C1 (Opal 570) | |
| | FISH Probe 3 | CD3e-C3 (Opal 620) | |
| | FISH Probe 4 | Abgre1-C4 (Opal 690) | |
| FISH Probe 1 | probe_contrast_threshold | 0.5 | Ncr1-C2 (Opal 520) |
| FISH Probe 2 | probe_contrast_threshold | 0.589 | Kcnj8-C1 (Opal 570) |
| FISH Probe 3 | probe_contrast_threshold | 0.477 | CD3e-C3 (Opal 620) |
| FISH Probe 4 | probe_contrast_threshold | 0.5 | Abgre1-C4 (Opal 690) |
| FISH Probe 1 | probe_min_intensity | 0.25 | Ncr1-C2 (Opal 520) |
| FISH Probe 2 | probe_min_intensity | 0.08 | Kcnj8-C1 (Opal 570) |
| FISH Probe 3 | probe_min_intensity | 0.199 | CD3e-C3 (Opal 620) |
| FISH Probe 4 | probe_min_intensity | 0.212 | Abgre1-C4 (Opal 690) |
| FISH Probe 1 | probe_min_spot_size | 0.5 | Ncr1-C2 (Opal 520) |
| FISH Probe 2 | probe_min_spot_size | 0.098214 | Kcnj8-C1 (Opal 570) |
| FISH Probe 3 | probe_min_spot_size | 0.2 | CD3e-C3 (Opal 620) |
| FISH Probe 4 | probe_min_spot_size | 0 | Abgre1-C4 (Opal 690) |
| FISH Probe 1 | probe_max_spot_size | 20 | Ncr1-C2 (Opal 520) |
| FISH Probe 2 | probe_max_spot_size | 20 | Kcnj8-C1 (Opal 570) |
| FISH Probe 3 | probe_max_spot_size | 20 | CD3e-C3 (Opal 620) |
| FISH Probe 4 | probe_max_spot_size | 20 | Abgre1-C4 (Opal 690) |
| FISH Probe 1 | probe_copy_intensity | 0.15 | Ncr1-C2 (Opal 520) |
| FISH Probe 2 | probe_copy_intensity | 0.15 | Kcnj8-C1 (Opal 570) |
| FISH Probe 3 | probe_copy_intensity | 0.15 | CD3e-C3 (Opal 620) |
| FISH Probe 4 | probe_copy_intensity | 0.15 | Abgre1-C4 (Opal 690) |
| FISH Probe 1 | probe_segg_agg | 0.95 | Ncr1-C2 (Opal 520) |
| FISH Probe 2 | probe_segg_agg | 0.95 | Kcnj8-C1 (Opal 570) |
| FISH Probe 3 | probe_segg_agg | 0.95 | CD3e-C3 (Opal 620) |
| FISH Probe 4 | probe_segg_agg | 0.95 | Abgre1-C4 (Opal 690) |

**Table S3**: Single cell RNAseq clustering of isolated mouse splenic cells

| Cluster | Number | Percentage | Cell type |
|---|---|---|---|
| 0 | 5126 | 32.3 | NK cells (NKp46+ NK1.1+ CD11b+ CD27low) |
| 1 | 4297 | 27.1 | NK cells (NKp46+ NK1.1+ CD11b+ CD27+) |
| 2 | 1341 | 8.4 | NK cells (NKp46+ NK1.1+ CD11b- CD27low) |
| 3 | 1227 | 7.7 | NK cells (NKp46+ NK1.1+ CD11b- CD27-) |
| 4 | 949 | 6.0 | B cells (Ebf1+) |
| 5 | 490 | 3.1 | Splenic red pulp macrophages (Hbb-bs+) |
| 6 | 454 | 2.9 | Splenic basophils (Tbc1d4) |
| 7 | 329 | 2.1 | Splenic dendritic cells (Cst3+ H2Ab1+) |
| 8 | 281 | 1.8 | Splenic eosinophils (Pf4+) |
| 9 | 250 | 1.6 | Splenic eosinophils (Dcn+) |
| 10 | 242 | 1.5 | Splenic germinal center centroblasts (Stmn1+ Top2a+) |
| 11 | 219 | 1.4 | CD8+ T cells (d8b1+) |
| 12 | 180 | 1.1 | Splenic basophils (Ifitm1+) |
| 13 | 175 | 1.1 | Splenic plasma cells (Igkc+) |
| 14 | 153 | 1.0 | Splenic NKT cells (Il7r+ Cd4+ Tnf+) |
| 15 | 136 | 0.9 | Splenic neutrophils (Lyz2+ Ifitm3+) |
| 16 | 29 | 0.2 | Neutrophils or eosinophils (S1009+ Retnlg+) |

Single cell RNAseq data analysis and clustering was performed using Seurat. Genes with expression enriched in the different clusters are shown in brackets. Cell cluster identification was performed by searching the ImmGen database and identifying the immune cell types that express the enriched gene(s).
